# Metabarcoding reveals the effect of land cover and temporal variations on the diet of Brown long-eared and Soprano pipistrelle bats within a unique European habitat, the pasture dominated landscapes of Ireland

**DOI:** 10.1101/2025.05.23.655448

**Authors:** Gwenaëlle Hurpy, Tina Aughney, Ilze Skujina, Niamh Roche, Emma C. Teeling

## Abstract

Considered as tertiary keystone predators, insectivorous bats play essential roles in maintaining the functioning of ecosystems. Investigating how bat species’ diets vary across landscapes is crucial for understanding bat ecology and their role in ecosystem health. Here, we characterised the predator-prey interactions of two common bat species with different foraging strategies, the Brown long-eared bat (*Plecotus auritus*) and Soprano pipistrelle (*Pipistrellus pygmaeus*), across the unique pastureland dominated landscape of Ireland. Over three years (2021-2023), faecal samples (n=4,627 in total) were collected annually at three time points (gestation, lactation, post-lactation) from twelve maternity roosts and analysed using metabarcoding and next-generation sequencing. Both bat species showed broad diet diversity, with 392 and 350 arthropod species identified for Brown long-eared bat and Soprano pipistrelle, respectively, primarily Lepidoptera and Diptera. The Brown long-eared bat exhibited a generalist diet demonstrating dietary flexibility. Lepidoptera interactions were more frequent overall (62%) compared to Diptera (31%), but interactions with Diptera species increased markedly at one specific roost, highlighting its capacity to adjust its diet to local prey availability. In contrast, the Soprano pipistrelle exhibited a more specialised diet, with 83% consisting of Diptera species. Both spatial and temporal factors significantly influenced dietary richness and composition in both species. Surrounding land cover, in particular, played an important role in shaping diet composition. Our findings suggest that the Brown long-eared bat exhibits a broad foraging strategy, acting as a generalist with a preference for Lepidoptera, while the Soprano pipistrelle shows a consistent reliance on Diptera. This study underscores the reliance of two widespread bat species, which play an important role in ecosystem well-being, on diverse insect taxa, highlighting the urgent need to conserve these insect populations.

## 1. Introduction

The landscape of Ireland is clearly distinct within Europe, with grasslands covering 57–61% of the land, the highest proportion in Europe compared to an average of 17.4% across Europe (Haughey, 2021; Eurostat, 2024). Forest cover remains low (11–14%) relative to the EU average (41.1%), and cropland accounts for just 5–9.5%, ranking Ireland among the lowest in Europe alongside Sweden and Finland (Eurostat, 2024). This unique landscape structure, dominated by grasslands and wetlands, suggests that Irish bat diets might differ from those in more forested or agricultural regions of Europe. Additionally, Ireland’s relatively low overall biodiversity may further shape prey availability and dietary patterns (McCarthy, 1986; Ferriss et al., 2009; Nelson et al., 2011; Kelly-Quinn and Regan, 2012; Harrison, 2014; Feeley et al., 2020).

Insectivorous bats feed on a broad diversity of arthropods, and contribute to well-functioning ecosystems (Kunz et al., 2011; Ghanem and Voigt, 2012; Lacher et al., 2019; Ramírez- Fráncel et al., 2022). Numerous studies have highlighted the importance of ecosystem services provided by insectivorous bats to agriculture, agroforestry, and even human health (Puig-Montserrat et al., 2020; Tiede et al., 2020; Charbonnier et al., 2021; Montauban et al, 2021; Da Silva et al., 2023; Hunninck et al., 2022). However, the majority of these studies focused on the services provided in relation to a single crop or one specific type of landscape, such as vineyards or forests. To inform national policy effectively, it is crucial to investigate the ecosystem services provided by bats on a broader scale (Braat and De Groot, 2012; Guerry et al., 2015). This requires a comprehensive understanding of bat diets across diverse and representative landscapes within countries.

To date, only five studies have investigated the diet of bats in Ireland, focusing on a limited number of species and sites (McAney et al., 1989; Shiel et al., 1991; Shiel et al., 1998; Flavin et al., 2001; Curran et al., 2022). Significant variations in the diet of the Lesser horseshoe bat (*Rhinolophus hipposideros*) (McAney et al., 1989) and Leisler’s bat (*Nyctalus leisleri*) (Shiel et al., 1998) were observed across roosts and seasons, demonstrating how environmental changes affect prey availability. Shiel et al. (1998) found that the diet of Leisler’s bats in Ireland showed lower richness compared to roosts in England and Germany, indicating that Irish land cover may significantly impact bat dietary diversity. These findings highlight that both landscape and temporal factors significantly influence bat diets. However, the limited studies underscore a considerable gap in understanding Irish bat feeding ecology.

Comprehensive research on bat diets across diverse habitats and timeframes is essential to understanding their ecological roles and ecosystem services, particularly as Ireland’s unique landscape may further shape the dietary habits of its local bat populations.

Among the nine bat species found in Ireland, the Soprano pipistrelle (*Pipistrellus pygmaeus*) is the second most abundant bat species (Roche et al., 2014). Although the Brown long-eared bat (*Plecotus auritus*) is less abundant, it is also widely distributed and regarded as common (Roche et al., 2014). Despite their prevalence, little is known about the diets of these two species in Ireland, presenting a knowledge gap in the understanding of their diet. The stability and accessibility of their maternity roosts, which are typically found in buildings from May to August, provide an ideal opportunity for non-invasive faecal sampling methods, making these species particularly suitable for diet studies. Furthermore, the distinct foraging behaviours of these species offer valuable insights into their ecological roles: the Brown long-eared bat is a gleaning species that forages within and on vegetation, while the Soprano pipistrelle uses aerial hawking to capture prey at the edges of vegetation (Swift, 1998; Roche et al., 2014; Dietz and Kiefer, 2016; Jones and Froidevaux, 2023). In terms of habitat, the Brown long- eared bat is typically associated with woodlands, including coniferous and broadleaf forests, but the species can also be found in riparian areas, treelines, scrubs, and orchards (Swift, 1998; Roche et al., 2014). In contrast, the Soprano pipistrelle is not always closely associated with woodlands, although it can be found in broadleaf woodlands. The species shows a particular preference for riparian habitats and low-density urban areas (Roche et al., 2014; Jones and Froidevaux, 2023).

Metabarcoding and next-generation sequencing techniques have revolutionised the investigation of diet composition across carnivorous, herbivorous, omnivorous, and insectivorous species (Shehzad et al., 2012; García-Estrada et al., 2012; Gill et al., 2023; Groen et al., 2023; Schumm et al., 2023; Tosa et al., 2023; Zurdo et al., 2023). These methods enable comprehensive dietary analysis, surpassing traditional studies that rely on morphological prey identification. This approach, now widely applied in bat studies (Galan et al., 2018; Alberdi et al., 2020; Tiede et al., 2020; Tournayre et al., 2021; Curran et al., 2022; Bourlat et al., 2023), provides comprehensive dietary profiles across numerous samples with high accuracy.

To provide a comprehensive understanding of the diets of the Brown long-eared bat and Soprano pipistrelle across Ireland, we collected faeces from multiple maternity roosts over three years at three sampling periods each year: gestation, lactation, and post-lactation. We used metabarcoding and NovaSeq Illumina sequencing to identify prey and bat DNA from these faeces. Specifically, we (1) examined the overall diet of the Brown long-eared bat and Soprano pipistrelle in Ireland, (2) compared the bat-prey interaction patterns between the Brown long-eared bat and Soprano pipistrelle and its variation among roosts, and (3) compared the richness and composition of their diets across different roosts, years, and months to identify the primary factors influencing dietary variation.

## 2. Materials and Methods

### 2.1 Bat faeces collection

#### 2.1.1 Identification of key bat roosts

Bat Conservation Ireland provided a list of 33 potential roosts for both Brown long-eared bat and Soprano pipistrelle. For each maternity roost, details were recorded, including building type, town, county, Irish grid coordinates, bat species present, feasibility of faecal sample collection (if available), and other relevant information. From this list, maternity roosts were pre-selected based on the following criteria: ease of access to faeces without disturbing the bats, likelihood of obtaining high-quality samples, and the characteristics of the surrounding landscape. A total of 19 maternity roosts–15 for Brown long-eared bats and four for Soprano pipistrelle–were sampled over a three-year period (2021 to 2023).

#### 2.1.2 Bat faeces sampling protocol

Bat faeces were collected during three sampling periods–gestation, lactation, and post- lactation–over a three-year period, from mid-May to the end of August in 2021, 2022, and 2023, from all pre-selected roosts. Sampling periods were chosen based on the life cycle of the two bat species (Aughney et al., 2018; Russo, 2023). All collections were approved by the Department of Housing, Local Government and Heritage of Ireland under license numbers DER/BAT 2021-39, DER/BAT-2022-48. Additionally, specific permits were obtained from the National Parks and Wildlife Service (NPWS) permits of Connemara and Wicklow Mountains National Parks. Faecal samples were collected using sterilised tweezers after old bat faeces had been removed and by placing newspaper sheets on the floor of bat maternity roost attics for two days (Puechmaille et al., 2007). All collected faeces were stored in individual tubes with silica beads to prevent DNA degradation (Taberlet et., al. 1999; Puechmaille et al., 2007), and were then stored at -20°C. Where indoor collection was not feasible, an outdoor collection system was established to gather droppings. Environmental conditions, including weather and wind, were assessed beforehand to optimise sample collection. Samples were collected immediately after sunrise each morning to minimise DNA degradation, again using sterilised tweezers and placed into individual tubes with silica beads.

Because non-target bat species were observed in some of the maternity roosts sampled, a protective measure was implemented to minimise cross-contamination: only isolated faecal samples were collected. Droppings touching each other were excluded from sampling to ensure the reliability of the DNA analysis. To be included in the study, a roost had to provide faecal samples at each of the three sampling periods annually. Consequently, only eight Brown long-eared bat roosts met this criterion for three years of sampling. For Soprano pipistrelle roosts, two were included in 2021 and sampled for all three years, with two additional roosts sampled in 2022 and 2023. Therefore, this study included data from a total of twelve bat maternity roosts (Fig. 1). A total of 4,627 faecal samples were collected over the three years, comprising 3,566 faeces from Brown long-eared bat roosts, and 1,061 faeces from Soprano pipistrelle roosts. (Supporting Information Table S1 presents details of the number of faecal samples collected for each roost).

**Figure 1.**
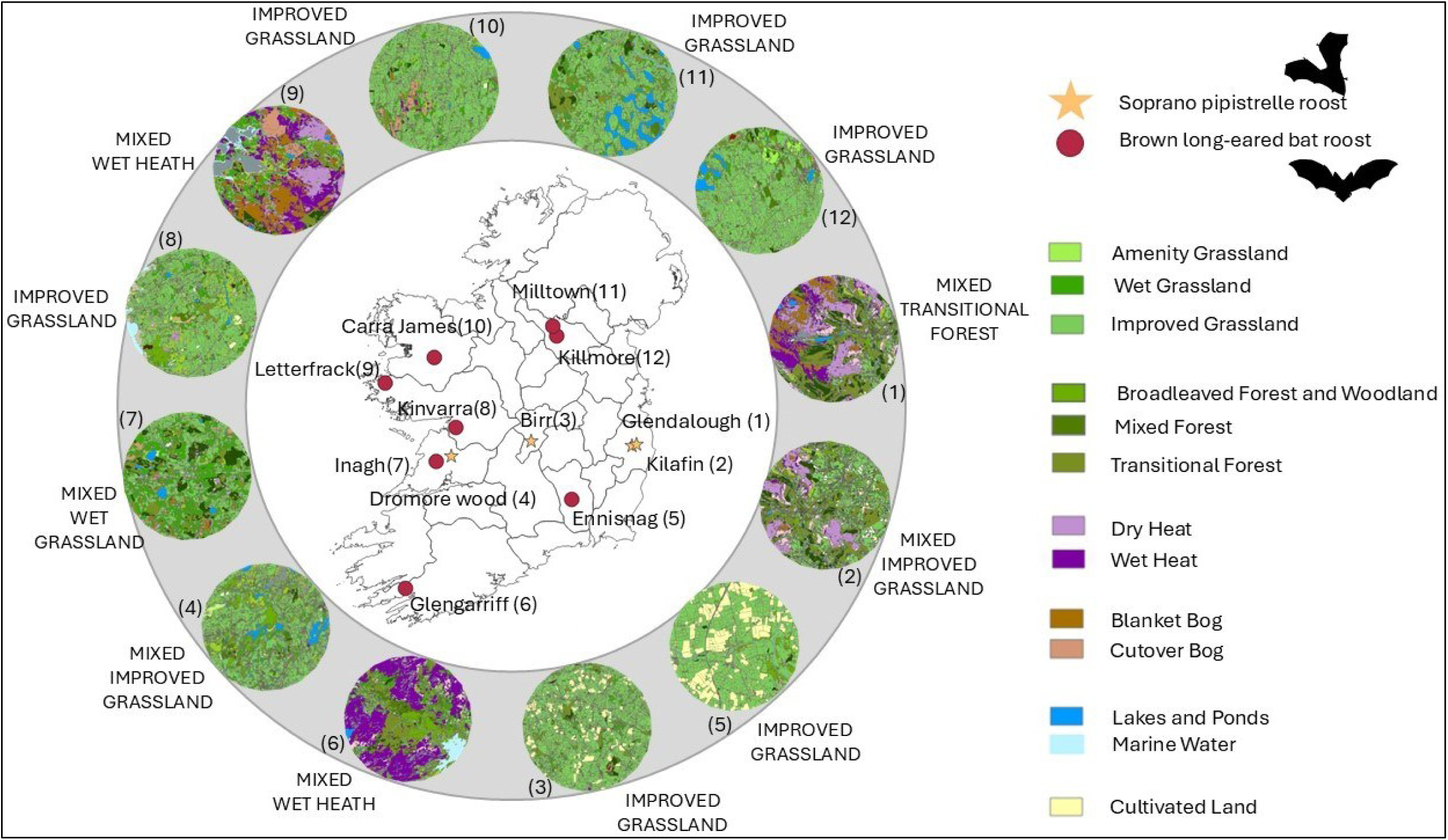
Location and main National Land Cover Map of Ireland of the roosts included in the analysis. 5km buffer for Soprano pipistrelle roosts: Glendalough (Co. Wicklow) (1), Kilafin (Co. Wicklow) (2), Birr (Co. Offaly) (3), Dromore Wood (Co. Clare) (4). 3km buffer for Brown long-eared bat roosts: Ennisnag (Co. Kilkenny) (5), Glengarriff (Co. Cork) (6), Inagh (Co. Clare) (7), Kinvarra (Co. Galway) (8), Letterfrack (Co. Galway) (9), Carra James (Co. Mayo) (10), Milltown (Co. Cavan) (11), Killmore (Co. Cavan)(12). Data land cover was extracted from the National Land Cover Map of Ireland (2023). National Land Cover Map of Ireland./ © Tailte Éireann - Surveying / Environmental Protection Agency.

### 2.2 Land cover data

The National Land Cover Map of Ireland (2023) was used to gather information on the landscape characteristics surrounding each maternity roost (Tailte Éireann Permit No. CYAL50442193). The distance of foraging sites from the roost varies between the two bat species. For the Brown long-eared bat, according to the literature, the foraging site distances range from a radius of 0 to 3.3 km from the roost (Entwistle et al., 1996; Swift, 1998; Dietz and Kiefer, 2018; Aughney et al., 2011; Ashrafi et al., 2013; Hillen et al., 2013). Therefore, we defined the potential foraging areas to be within a 3km radius from the roost. In contrast, for the Soprano pipistrelle, the range is broader, varying from 480 m to 12.3 km from the roost (Davidson-Watts and Jones, 2006; Nicholls and Racey, 2006; Stone et al., 2015; Dietz and Kiefer, 2018). Within Ireland, Bat Conservation Ireland has typically recorded Soprano pipistrelles foraging 5km from their roost (Tina Aughney pers.com). Therefore, we estimated the foraging area to be within a 5km radius. Land cover data were extracted using *QGIS Desktop 3.36.0* for buffers of 3 km and 5 km around the respective roosts of the two species (Fig. 1)

### 2.3 Preparation of sequencing libraries

DNA was extracted and sequenced from a maximum of 20 to 25 faecal samples per roost and sampling event, when available. DNA extraction was conducted annually after each fieldwork season. To avoid any cross-contamination across roosts, DNA extractions were processed by roost in 96-well plates. Arthropod and bat DNA were extracted following Zarzoso-Lacoste’s protocol (2018) with modifications (Supplementary Information Table S2). To minimise cross- contamination during the extraction process, all extractions were conducted in a fume hood, with UV sterilisation for 45 minutes before and after each extraction batch, and all reusable material was washed with a 10% bleach solution. Two primer sets targeting mitochondrial DNA Cytochrome c oxidase subunit I (COI) (Folmer et al., 1994; Hebert et al., 2003) were used to identify bat and prey species from extracted DNA. These primer sets, fwhF1/fwhR1 (Vamos et al., 2017) and ZBJ-ArtF1c/ZBJ-ArtR2c (Zeale et al., 2011) amplify partial COI regions of 178 bp and 157 bp, respectively. Tagged PCR was performed (Bohmann et al., 2022) with forward and reverse primers tagged with 8 bp unique sequences to allow for multiplexing of the faecal samples, following Taberlet (2018) and Illumina recommendations. Each 96-well plate included one negative extraction control, one negative PCR control, and one positive PCR control. To minimise contamination, PCRs were conducted in a different laminar fume hood to that used for the DNA extractions, which was also UV-sterilised for 45 minutes before and after PCR plate. For each PCR amplicon, the concentration was estimated using the QUBIT Hight sensitivity DNA protocol. PCR amplicons were pooled in equimolar concentrations into sequencing libraries by roost and year of sampling, generating a total of 74 libraries over the three years of the study. Pooled libraries were cleaned up using the AMPure XP Beads protocol (Beekman coulter).

The libraries were sent to Biomarker Technologies for an Illumina NovaSeq sequencing on NovaSeq 6000 platform, using 150 bp paired-end runs. Illumina adapters and indexes were added using ligation-based library preparation to minimise tag-jumping (Bohmann et al., 2022).

### 2.4 Bioinformatic – Recovery Bat and prey sequences

The identification of bat and prey species from NovaSeq sequences was conducted using the pipeline proposed by Browett et al. (2021) in combination with OBITools v1.2.13 (Boyer et al., 2016), VSEARCH (Rognes et al., 2016), and BLAST v2.12.0 (Camacho *et al*., 2009) scripts.

Additionally, R scripts from the Metabarpark Project (Wangensteen and Turon, 2017) were used to adapt file formats between successive steps. Each library was processed separately. Briefly, raw sequences from each library were ‘paired-ended’ using ‘*illuminapairedend*’ with a minimum score of 40. The next step was ‘*ngsfilter*’ used to demultiplex the sequences based on 8 bp sample indices and attribute them to the corresponding faecal samples. Sequences with unexpected lengths, those containing “N” base, or those occurring less than 10 times were removed. Additionally, chimeric sequences were identified using the ‘*uchimeout’* function. The sequences that remained after filtering, representing Molecular Operational Taxonomic Units (MOTUs), were clustered with 98% identity using ‘*sumaclust*’ command.

MOTUs were assigned to taxonomic identifiers (taxid), using remote BLAST against the National Center for Biotechnology Information (NCBI) database with ten sequences target by MOTUs (Altschul et al., 1990; Sayers et al., 2022), with a 98% identity and 90% coverage threshold (Tournayre et al., 2021). Where MOTUs were assigned to several species, the most recent common ancestor was selected (Wangensteen et al., 2018). The complete script is available in Supporting Information Bioinformatic Script.

To minimise potential false positives resulting from cross-contamination, MOTUs present in extraction or PCR blanks were removed from the library if their percentage of reads in blanks exceeded 10% compared to other samples (Wangensteen et al., 2018; Vescera et al., 2024). Additionally, a stringent threshold was applied to eliminate potential artifact sequences introduced during PCR and sequencing. MOTUs with occurrence levels below 1% in each faecal sample were removed (Mata et al., 2021) using the ‘*hill_div’* function from the *hilldiv* R package (version 1.5.1, Alberdi and Gilbert, 2019). The taxonomy of each MOTU was retrieved using the Taxonomizr R package v0.10.6 (Sherrill-Mix, 2023).

The third step involved identifying the sequence belonging either to, “host” (bat species), “prey” (species considered as prey), “external parasite” (arthropod species determined as a parasite of the bat species) and “other” (any other categories of species). The presence or absence in Ireland of each identified species taxa was verified using the National Biodiversity Data Center database (National Biodiversity Data Center, 2024) and scientific literature (O’Connor et al., 2009; Ashe et al., 2012; O’Connor and Nelson, 2012; Baker et al., 20213; Chandler 2024). Only species documented as present in Ireland in at least one of these databases were included for the final analysis. The current known distribution of prey species not recorded in Ireland was verified using Global Biodiversity Information Facility (GBIF) database (GBIF, 2024).

Finally, during sampling, multiple bat species were observed sharing roosts, and eventually, non-target bat species were observed urinating on newspapers placed to collect faecal samples, potentially introducing their DNA on faecal samples. DNA from urine was considered acceptable for analysis because it did not affect diet analysis results. However, a threshold was set to ensure analysis integrity. Only faecal samples containing at least 90% DNA from a single bat species were included, balancing data reliability with sample retention.

### 2.5 Statistical analysis

Statistical analyses were conducted using R (version 4.3.3) and R studio (version 2023.12.1.402). The *ggplot2* package (version 3.5.1 ; Wickham, 2016) was used for data visualisation. Following recommendations by Galan et al. (2018), PCR products with less than 500 reads were discarded. Faecal samples from each roost and sampling event were sequenced, with the number of samples varying by event. For sampling events with more than 20 faecal samples, a random selection of up to 20 samples was made using the ‘*sample*’ function to ensure unbiased representation.

To evaluate the reliability of the data, *‘iNEXT’* and ‘*ggiNEXT*’ functions of the *iNEXT* package (version 3.0.1) were used to generate sample completeness curves for each sampling event (Chao et al., 2014; Hsieh et al., 2016). These curves estimate sample coverage, which quantifies the proportion of the total species diversity in a community that is represented in the sample.

Prey richness in each faecal sample was calculated using the ‘*hill_div*’ function from the *hilldiv* package (version 1.5.1), with q set to 0. To compare prey richness between the Brown long- eared bat and Soprano pipistrelle, we performed a Wilcoxon-Mann-Whitney test (Wilcoxon, 1947 ; Mann and Whitney, 1947). Additionally, we used the Kruskal-Wallis test to assess differences in prey richness across roosts, sampling time points, and years (Nwobi and Akanno, 2021).

To explore the specificity of foraging patterns among bat maternity roosts, we assessed diet distinctiveness (DD) between roost as the one minus the pairwise Pianka’s niche overlap (Pianka, 1973; Roswag et al., 2018; Mata et al., 2021), using the ‘*niche.overlap’* function from the *spaa* package (version 0.2.2, Zhang, 2016). Pairwise Pianka’s niche overlap values were verified using the bootstrap function ‘*niche.overlap.boot*’ function from the *spaa* package. To visualise interaction patterns between prey arthropod order and maternity roosts, we then generated web plots using the ‘*bipartiteD3*’ function from the *bipartiteD3* package (version 0.3.2, Terry, 2024) for the two bat species.

To investigate the effects of spatial (maternity roost location) and temporal (year and sampling event) factors on prey composition and dispersion, we first created a distance matrix using the ‘*vegdist*’ function with the Jaccard method based on the presence/absence of prey from the *vegan* package (version 2.6.6, Oksanen et al., 2024). We then performed a permutational multivariate analysis of variance (PERMANOVA) to assess differences in beta diversity (i.e. variation in prey composition) among roosts, years, and events, applying the ‘*adonis2* ‘ function with 999 permutations (Anderson, 2006; Anderson et al., 2006; Anderson, 2014). To visualise compositional patterns between bat species and roost, we generated a non-metric multidimensional scaling (nMDS) plot using the ‘*metaMDS*’ function (Tong, 1992; Zuur et al., 2007). To assess whether roost or year had a stronger influence on dietary composition, we constructed a dendrogram using Jaccard dissimilarity and complete linkage clustering with the ‘*hclust’* function from the *stats* package (v3.6.2). Finally, to examine the relationship between prey composition and landscape diversity, we performed a Mantel test based on Spearman correlation using the ‘*mantel’* function (Borcard and Legendre, 2012; Tournayre et al., 2021).

## 3. Results

### 3.1 Landscape characteristics

According to the National Land Cover Map of Ireland, both Brown long-eared bat and Soprano pipistrelle maternity roosts were located within a mosaic of natural or semi-natural habitats. Five Brown long-eared roosts were located in areas where “Improved grassland” accounted for more than 50% of the land cover within a 3 km radius. The remaining three roosts were located in areas characterised by a variety of land cover types, none of which individually represented more than 50% of the surrounding area. These diverse land cover types included “Wet heath”, “Dry heath”, “Broadleaved Forest”, Transitional Forest”,

“Improved grassland”, “Wet grassland”, “Coniferous Forest”, and “Blanket bog” (Fig. 1). In contrast, three of the Soprano pipistrelle roosts were located within a mix of land cover types, each contributing less than 50% of the land within the 5 km radius. These land cover types included “Transitional Forest”, “Dry heath”, “Wet heath”, “Coniferous Forest”, ”Improved grassland”, “Broadleaved Forest”, and “Scrub”. Only one roost was located in an area dominated by one land cover type, with “Improved grassland” accounting for 67.2% of the surrounding area (Fig. 1). The percentage of each landcover levels 1 and 2 are available in Supporting Information Data Set S1.

### 3.2 Prey and bat species identification

The 74 NovaSeq sequencing runs generated a total of 476,391,107 reads, with individual runs producing between 3,903,247 and 14,480,797 reads. Raw data and Metadata are available on DataDryad (DOI: https://doi.org/10.5061/dryad.qbzkh18vt). After applying bioinformatic filtering, 255,022,827 reads were retained for further analysis. After filtering out MOTUs with less than 1% occurrence across faecal samples, we identified 908 taxa at the species level. Each species was assigned to a category (prey, bat host, external parasites, or other), resulting in a total of 691 prey species identified for both the Brown long-eared bat and Soprano pipistrelle. Of these, 51 species were not previously recorded in Ireland and were excluded from the final analysis. The overall list of the 907 identified species, along with the 51 non-Irish species and their known distributions, is provided in Supporting Information Data Set S2.

In terms of host species, five bat species were identified: Brown long-eared bat (1,257 faeces), Soprano pipistrelle (475 faeces), Lesser horseshoe bat (*Rhinolophus hipposideros*) (168 faeces), Natterer’s bat (*Myotis nattereri*) (44 faeces), and Common pipistrelle (*Pipistrelle pipistrellus*) (14 faeces). Brown long-eared bat was identified as the host with 100% confidence for 1163 faeces (93% of the faecal samples). For faecal samples containing DNA from multiple bat species, 53% were identified as containing Brown long-eared DNA at 98% or 99% confidence. Similarly, Soprano pipistrelle was identified as the host in 460 faeces (97%) with 100% confidence. Among the faecal samples containing DNA from other bat species, 41% showed Soprano pipistrelle DNA at 98% or 99% confidence.

Sampling coverage was estimated across all sites and years, using the iNEXT and ggiNEXT functions. All roosts exceeded 91%, indicating a high level of data reliability. However, when coverage was assessed at each roost for individual sampling event, variability was observed with some sampling periods showing lower coverage (Supplementary Information Figure S1).

### 3.3 Factors influencing prey richness in the diet

A total of 16 orders were identified in the diets of the Brown long-eared bat and Soprano pipistrelle, with 14 and 11 arthropod orders, respectively. Lepidoptera and Diptera species were the most prevalent prey for Brown long-eared bat, with 184 and 154 species, respectively. The diet of Soprano pipistrelle was largely dominated by Diptera species (210 species), while 83 species of Lepidoptera were recorded (Fig. 2). Prey richness per faecal sample ranged from 1 to 21 species for the Brown long-eared bat and from 1 to 20 for the Soprano pipistrelle, with an average of 6.3 and 7.06 species, respectively (Fig. 3). This difference in prey richness between the two species was statistically significant (p = 3.483e- 16; Table 1). Within both bat species, prey richness varied significantly across the roosts and from year to year (Fig. 3). Although prey richness differed across years, there was no significant difference between sampling periods (Table 1).

**Figure 2.**
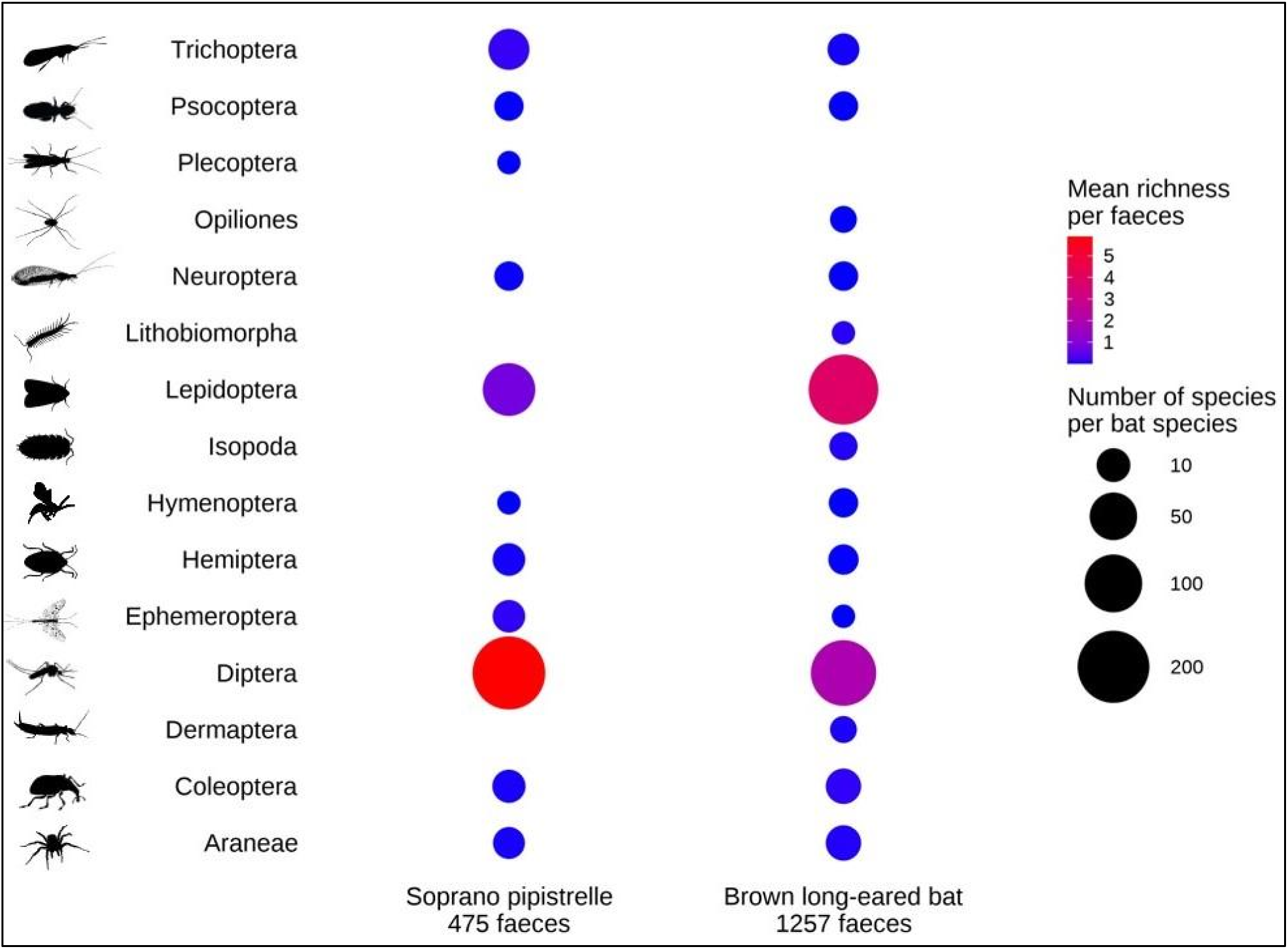
Dot plot showing total prey species by order in the diet of Brown long-eared bat and Soprano pipistrelle along with the mean species per order.

**Figure 3.**
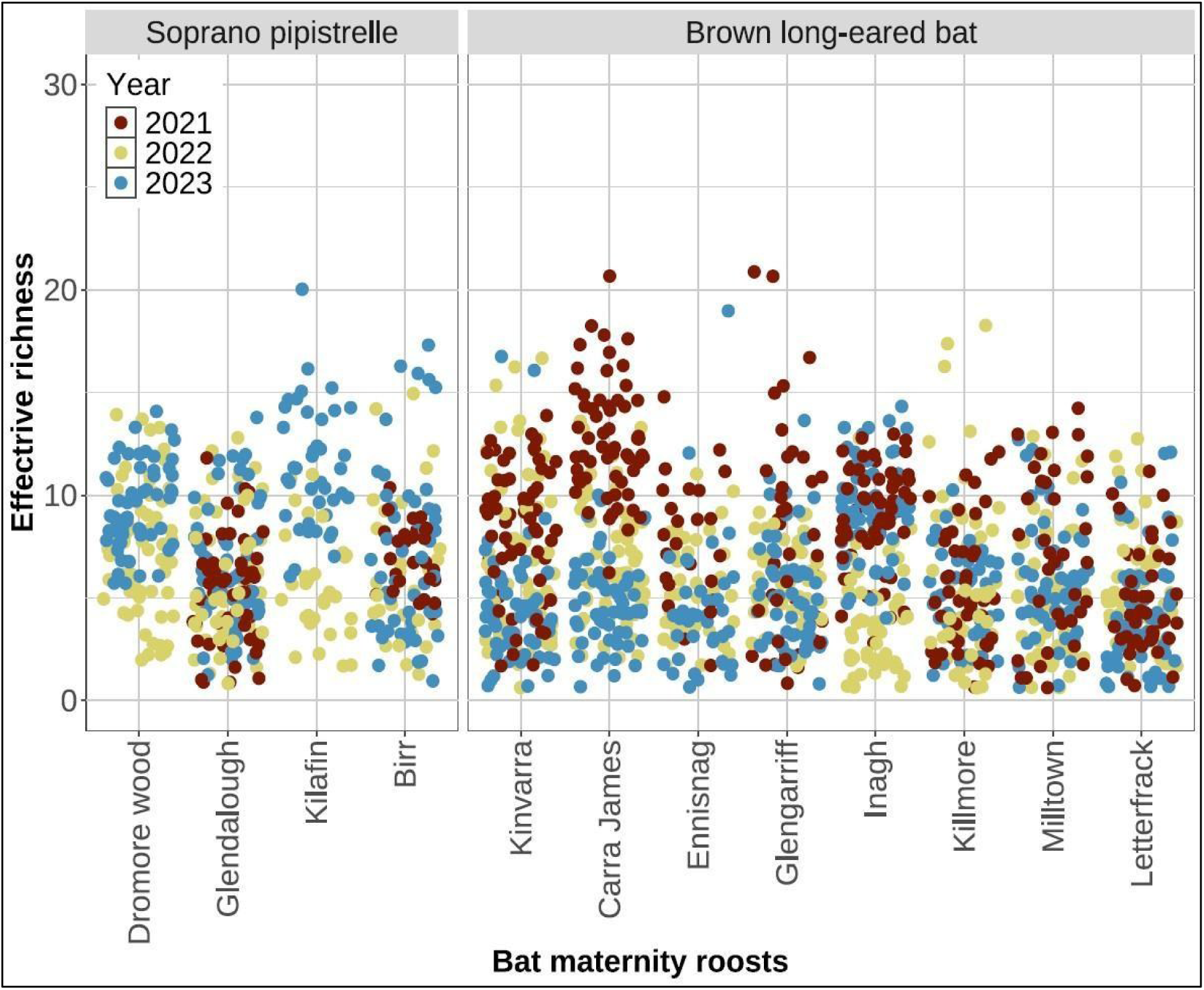
Jitter plot showing the effective richness for each faecal sampled analysed per bat species and bat maternity roost. Colours represent the year of sampling collection. Effective richness represents the number of prey species per faecal samples.

**Table 1.** Results of comparison Kruskal Wallis tests for comparing prey species richness for each variable: bat species, roost, year, and sampling period. Significant p-value (p-value < 0.05) are indicated in bold.

| Bat species | Variable tested | Kruskal Wallis test results |  |  |
| --- | --- | --- | --- | --- |
|  |  | X<sup>2</sup> | Df | P-value |
| Brown long-eared bat | Year | 112.67 | 2 | **< 2.2E-16** |
|  | Roost | 105.73 | 7 | **< 2.2E-16** |
|  | Sampling period | 4.22 | 2 | 0.121 |
| Soprano pipistrelle | Year | 47.16 | 2 | **5.76E-11** |
|  | Roost | 59.27 | 3 | **8.41E-13** |
|  | Sampling period | 0.52 | 2 | 0.771 |
| Brown long-eared bat<br>and<br>Soprano pipistrelle | Bat species | 21.53 | 1 | **3.48E-06** |

The mean richness of arthropod orders varied across the sampling period and roost for both bat species. In the diet of the Brown long-eared bat, richness of the two dominant orders– Lepidoptera and Diptera– showed variation across both roosts and sampling time points, with no single arthropod order consistently dominating. Lepidoptera richness showed strong variation across roosts, with a mean ranging from 0.27 to 12.00 and a maximum richness up to 17 species while Diptera order presented a mean richness ranging from 0.15 to 6.00. In contrast, in Soprano pipistrelle roosts, Diptera species consistently had the highest prey richness, with mean values per roost ranging from 2.80 to 10.45, and a maximum richness up to 17 species. Lepidoptera, the second dominant order, showed lower mean richness, ranging from 0.05 to 4.50, with a maximum of seven in one of the roosts sampled. Other arthropod orders showed low mean richness across all roosts, all below 1.00 (Supplementary Figure S2).

### 3.4 Bat and prey interactions

DD values range from 0 (low dietary distinctiveness) to 1 (high distinctiveness). DD between the two bat species was 0.82, indicating limited overlap in prey composition. Pianka’s niche overlap value was 0.172, with a bootstrap estimated of 0.182, providing confidence in this result. Web plot analysis revealed that the Soprano pipistrelle’s diet was primarily dominated by Diptera, which accounted for 83.9% of prey interactions. In contrast, the Brown long-eared bat’s diet was more balanced, with Lepidoptera (61.66%) and Diptera (31.09%) being both dominant prey orders (Fig. 4).

**Figure 4.**
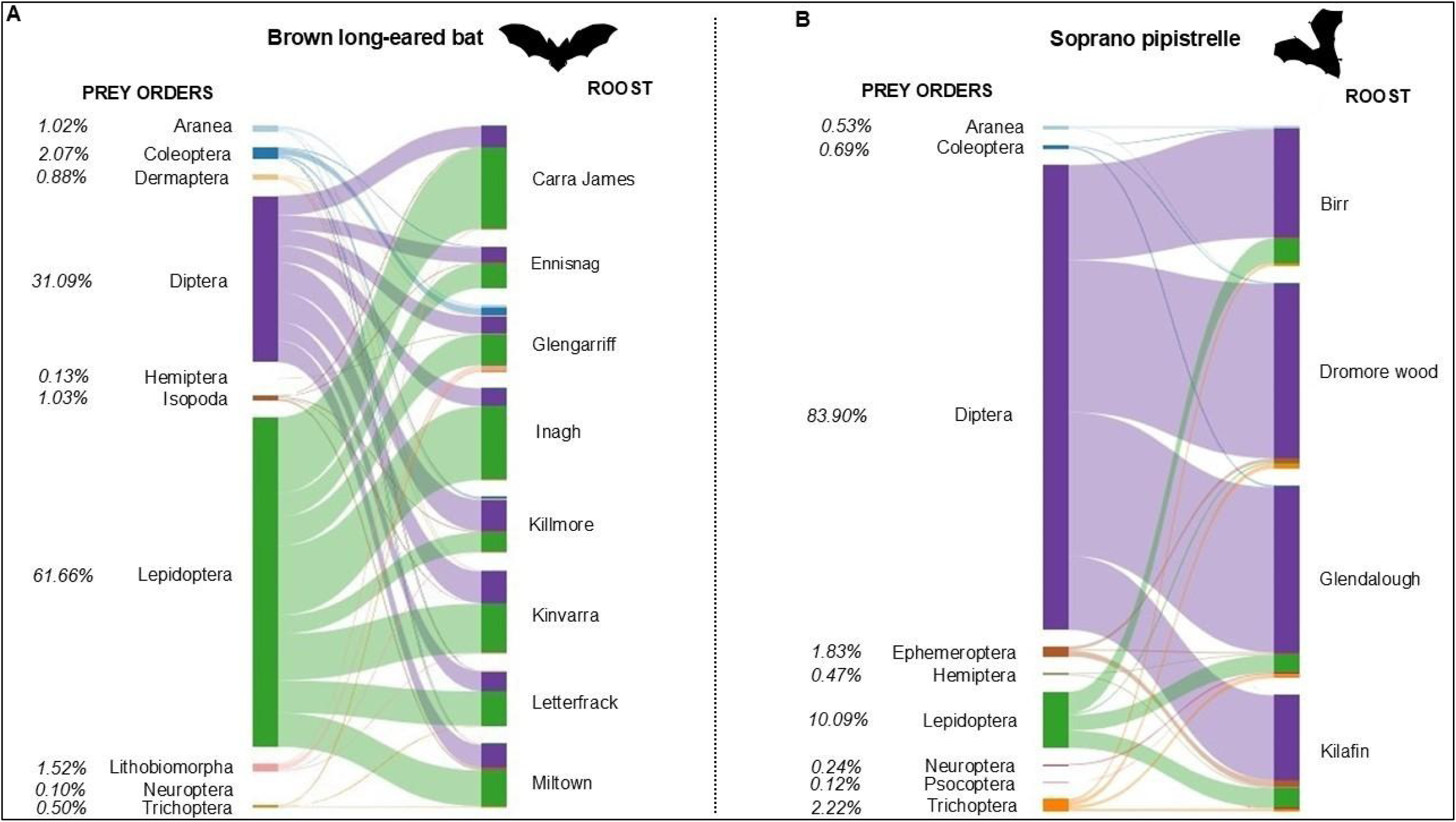
Web plots showing interactions between prey orders with Brown long-eared bat (A) and Soprano pipistrelle (B) across maternity roosts. Percentages show the percentage of each order in diets. Only orders representing more than 0.1% of the total interactions are presented. The web plot was created using the ‘bipartiteD3’ function from the bipartiteD3 package.

At the family level, both bat species interacted with a wide range of Lepidoptera and Diptera families. The Brown long-eared bat interacted with 21 Lepidoptera and 29 Diptera families, while the Soprano pipistrelle interacted with 21 Lepidoptera and 35 Diptera families. However, most of these families contributed only a small proportion to the overall interactions. Among the Brown long-eared bat’s interactions, the most frequently encountered Lepidoptera families were Noctuidae (68.50%), Hepialidae (9.66%), Geometridae (8.93%), and Erebidae (8.58%), while 17 other families each contributed less than 5%. Within Diptera, Muscidae (13.38%) and Scathophagidae (9.83%) were present, but Tipulidae (57.06%) was dominant, representing the majority of interactions. The remaining 26 Diptera families contributed less than 5% each. In the Soprano pipistrelle’s diet, the primary Lepidoptera families were Blastobasidae (28.15%), Hepialidae (16.13%), Noctuidae (13.49%), Tortricidae (12.02%), Geometridae (7.04%), and Gracillariidae (6.45%), while 15 additional Lepidoptera families contributed less than 5% each. Among Diptera, the dominant families were Psychodidae (23.28%), Chironomidae (22.96%), Limoniidae (14.81%), and Tipulidae (7.13%), with 29 other families each representing less than 5%. Detailed information on the contribution of each family to the diet is provided in Supporting Information Figure S3.

Within species, DD values between roosts ranged from 0.258 to 0.740 for the Brown long- eared bat, and from 0.400 to 0.662 for the Soprano pipistrelle. Most interspecific roost comparisons showed high DD values, with the exception of one Soprano pipistrelle roost (Birr), located in an improved grassland landscape. This roost exhibited moderate dietary dissimilarity with several Brown long-eared bat roosts, with DD values ranging from 0.557 (Ennisnag; improved grassland) to 0.857 (Glengarriff; mixed wet heath and woodland) (Table 2)The Brown long-eared bat exhibited variability in interactions across roosts, with Lepidoptera and Diptera contributing between 35.10% to 80.02% and 19.29% to 55.18% of the total of interactions with prey species, respectively. At the Glengarriff roost, three additional arthropod orders each accounted for more than 5% of the interactions: Araneae (5.10%), Coleoptera (10.51%), and Lithobiomorpha (8.17%). Similarly, the Soprano pipistrelle’s interactions with Lepidoptera and Diptera varied across roosts, ranging from 1.01% to 17.63% and 72.86% to 93.64%, respectively. However, Diptera consistently dominated the diet, accounting for more than 70% of prey interactions at all roosts.

**Table 2.** Diet distinctiveness values between all roosts included in the studies, Brown long-eared (in black) and Soprano pipistrelle (in blue). Diet Distinctiveness between each roost was estimated as minus one the Pairwise Pianka’s niche overlap value.

|  | **Dromore**<br>**wood** | **Glendalough** | **Kinvarra** | **Carra**<br>**James** | **Ennisnag** | **Miltown** | **Glengarriff** | **Letterfrack** | **Inagh** | **Killmore** | **Birr** |
| --- | --- | --- | --- | --- | --- | --- | --- | --- | --- | --- | --- |
| **Glendalough** | 0.400 |  |  |  |  |  |  |  |  |  |  |
| **Kinvarra** | 0.955 | 0.966 |  |  |  |  |  |  |  |  |  |
| **Carra James** | 0.958 | 0.958 | 0.302 |  |  |  |  |  |  |  |  |
| **Ennisnag** | 0.946 | 0.944 | 0.343 | 0.395 |  |  |  |  |  |  |  |
| **Miltown** | 0.881 | 0.864 | 0.506 | 0.343 | 0.493 |  |  |  |  |  |  |
| **Glengarriff** | 0.974 | 0.947 | 0.614 | 0.567 | 0.707 | 0.562 |  |  |  |  |  |
| **Letterfrack** | 0.977 | 0.942 | 0.740 | 0.483 | 0.657 | 0.522 | 0.606 |  |  |  |  |
| **Inagh** | 0.976 | 0.972 | 0.506 | 0.258 | 0.548 | 0.408 | 0.631 | 0.414 |  |  |  |
| **Killmore** | 0.926 | 0.903 | 0.422 | 0.475 | 0.367 | 0.404 | 0.625 | 0.616 | 0.550 |  |  |
| **Birr** | 0.657 | 0.678 | 0.690 | 0.751 | 0.557 | 0.619 | 0.827 | 0.787 | 0.852 | 0.576 |  |
| **Kilafin** | 0.662 | 0.481 | 0.928 | 0.981 | 0.836 | 0.906 | 0.936 | 0.913 | 0.972 | 0.830 | 0.478 |

Ephemeroptera contributed more than 5% of the diet at one roost, Kilafin, representing 5.82% of the total interactions (Fig. 4).

### 3.5 External factors influencing beta diversity

Initial nMDS analysis revealed the presence of outliers, which were subsequently removed to ensure a more robust and accurate assessment of beta diversity. The revised results showed a better fit of the ordination, indicating improved clarity in the separation of prey compositions between bat species and roosts. The list of outliers is available in Supporting Information Table S3. Composition of diet significantly varied depending on bat species and spatial and temporal factors. PERMANOVA results revealed that the prey composition varied significantly between bat species, bat roosts, sampling periods and year (Table 3). The nMDS plot showed clear separation between bat species, but with variability across roosts within bat species. The stress value of 0.08, indicating a good fit for the ordination (Fig. 5 and Supporting Information Figure S4).

**Figure 5.**
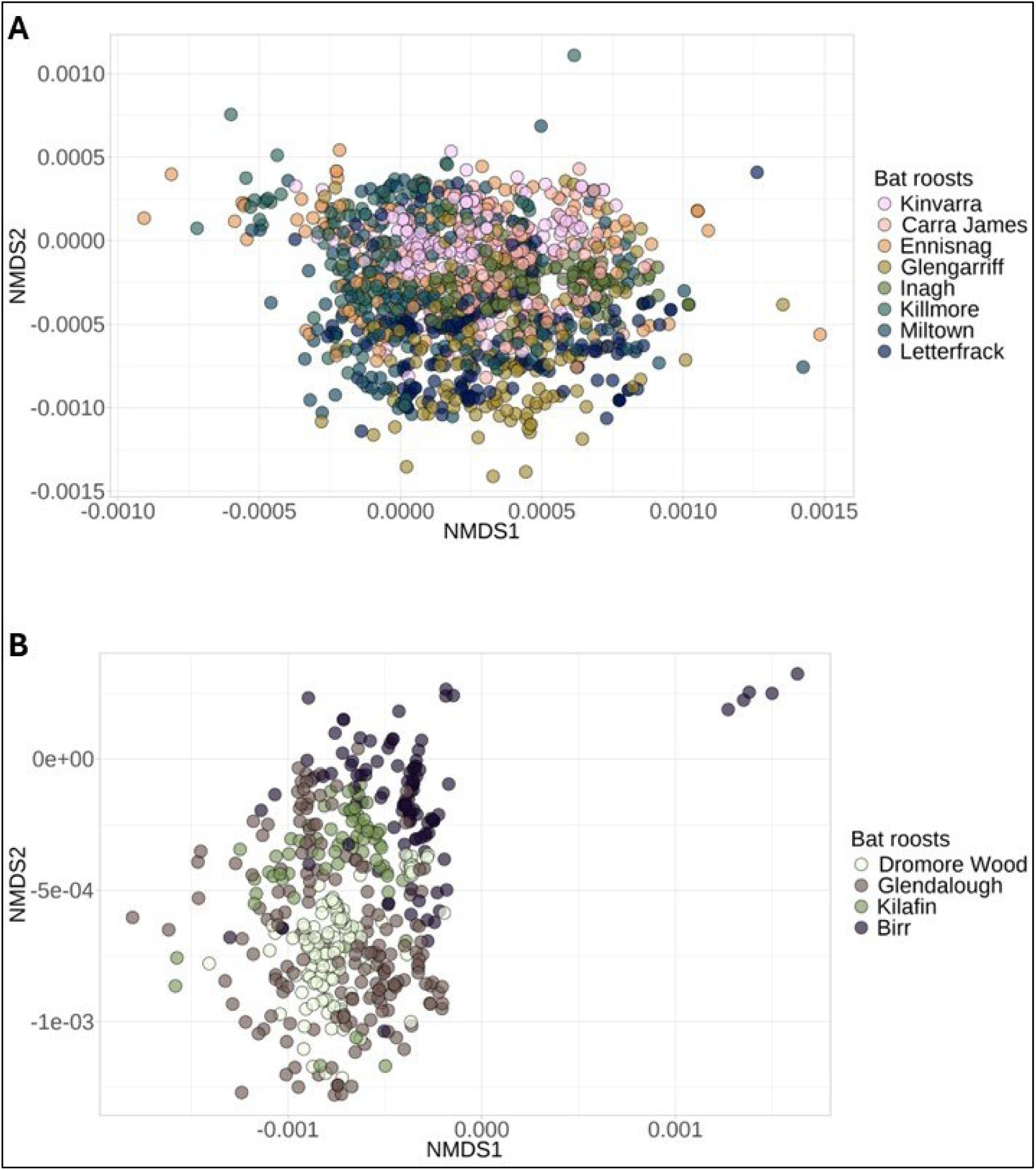
Non-metric multidimensional scaling plot showing the dissimilarity in diet composition across bat maternity roosts of Brown long-eared bat (A) and Soprano pipistrelle (B). Each point represents, a faecal sample, and the distance between points reflects the similarity of species composition based on Jaccard dissimilarity. The stress value of 0.08 indicates a good representation of the data in two dimensions. Dissimilarity matrix used to create the plot was performed using ‘vegdist’ function of Vegan package. Plot was created with ‘metaMDS’ function. For better readability two samples were taken out the plot in (A), the original plot is available in Supplementary Information S4.

**Table 3.** Results of PERMANOVA (A) and MANTEL tests (B), used to identify significant differences in diet composition between bat species, maternity roosts, year of sampling, and sampling period. Table presents F-statistics, R^2^ values, degree of freedom, and p-values (Pr(>F). PERMANOVA test (performed using ‘adonis2’ function of vegan package) compares diet composition between groups. MANTEL test measures correlation between beta diversity and landscape diversity, using ‘mantel’ with spearman correlation. MANTEL results are indicated with 95% CI [0.50, 0.75]. Significant p-values (< 0.05) are indicated in bold.

| (A) | **PERMANOVA based on 999 permutations** |  |  |  |
| --- | --- | --- | --- | --- |
|  | Df | R<sup>2</sup> | F | Pr(>F) |
| Bat species | 1 | 0.0563 | 102.47 | 0.001 |
| Roost | 11 | 0.1866 | 35.621 | 0.001 |
| Year | 1 | 0.0118 | 20.582 | 0.001 |
| Sampling period | 2 | 0.0192 | 16.881 | 0.001 |
| Year * Sampling period | 8 | 0.0486 | 10.99 | 0.001 |

| (B) | **MANTEL TEST based on 999 permutations** |  |  |  |  |  |
| --- | --- | --- | --- | --- | --- | --- |
|  | Statistic r | Significance | 90% | 95% | 97.50% | 99% |
| Brown long-eared bat | 0.2525 | 0.001 | 0.0107 | 0.0143 | 0.0165 | 0.0190 |
| Soprano pipistrelle | 0.3161 | 0.001 | 0.0060 | 0.0082 | 0.0093 | 0.0115 |

Finally, hierarchical clustering of diet composition was performed for the eight Brown long- eared bat and four Soprano pipistrelle roosts from 2021 to 2023 (Fig. 6), using the complete linkage method with Jaccard presence/absence similarity. The analysis showed that the diet compositions of Brown long-eared bat roosts from 2021 were more similar to each other compared to those from 2022 and 2023. Additionally, the results indicated that diet similarity between years varied depending on the roost. For the Soprano pipistrelle roost in Birr, the clustering suggested greater similarity in diet composition across years compared to other roosts investigated.

**Figure 6.**
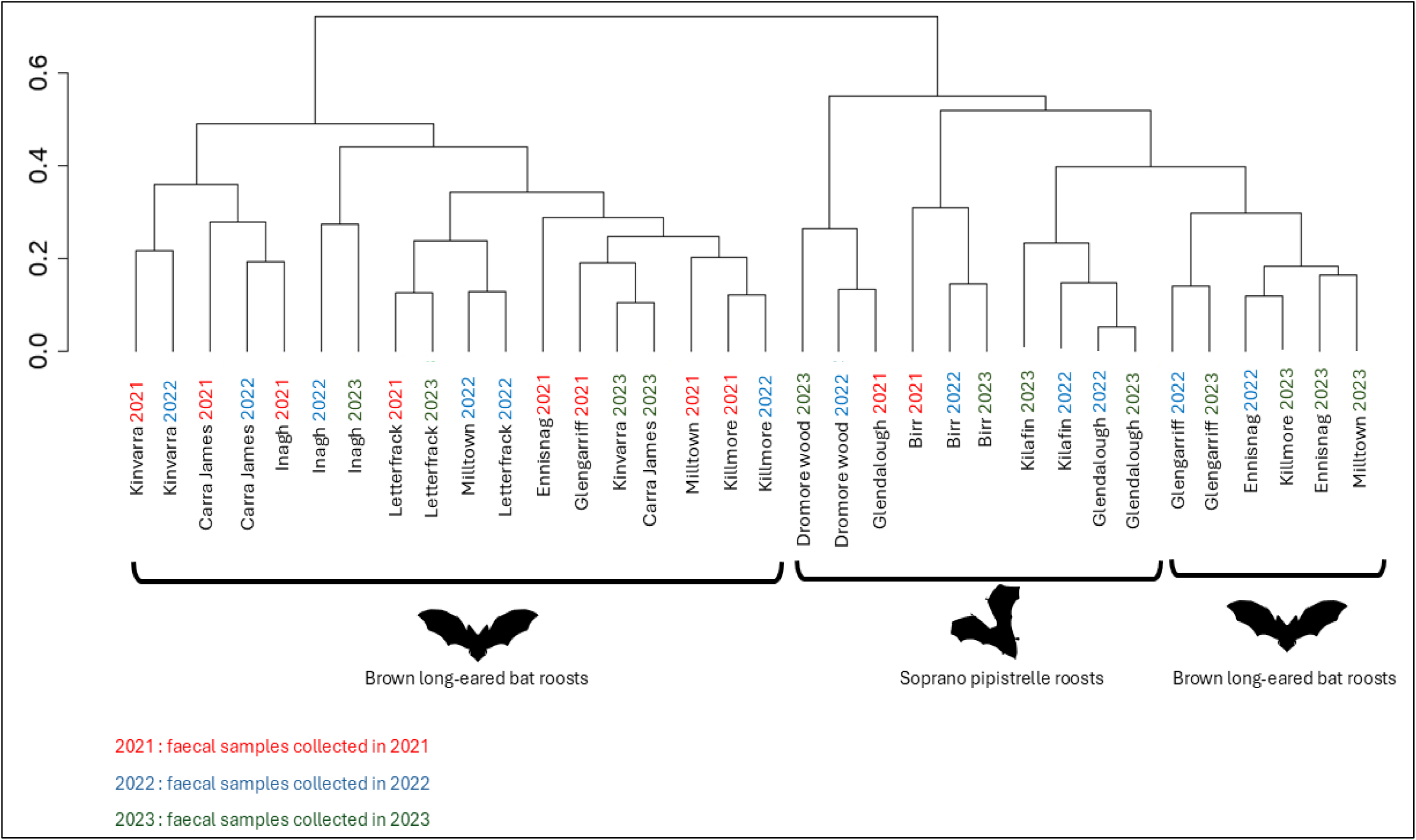
Dendrogram of hierarchical clustering of diet composition across the twelve bat maternity roosts from 2021 to 2023. Clustering was performed using Jaccard dissimilarity and complete linkage method, with the ‘hclust’ function of the stats package.

Mantel tests revealed a significant positive but low correlation between the dissimilarity of the beta diversity and the dissimilarity of the land cover among roosts for each bat species, with a higher correlation for the Soprano pipistrelle roosts (Table 3). This indicated that the more the landscape surrounding the roost differs, the more the prey composition differs.

## 4. Discussion

The dominance of grassland, coupled with low levels of forest and arable land makes the Irish landscape unique within Europe (Haughey et al., 2021; Eurostat, 2024). In this study, we used DNA metabarcoding to analyse the diet of two common bat species: the Brown long- eared bat and the Soprano pipistrelle. Over a three-year period, we collected faecal samples from twelve maternity roosts–eight Brown long-eared bat roosts and four Soprano pipistrelle roosts–using a non-invasive sampling protocol across various landscape types that represent the diverse land cover of Ireland.

### 4.1 Brown long eared bat’s diet

The Brown long-eared bat revealed a diverse diet, with 392 taxa identified across 15 arthropod orders. Lepidoptera was the dominant prey group, comprising 184 species and accounting for 62% of interactions, followed by Diptera with 154 species which contributing to 31% of interactions. This diversity of arthropods, with both flying and non-flying prey, such as Aranea, Opiliones, and Lithobiomorpha is consistent with other metabarcoding studies (Razgour et al., 2011; Andriollo et al., 2019). As Razgour et al. (2011) and Rostovskaya et al. (2000), Noctuidae species were highly represented in the diet confirming the high ability of the Brown long-eared bat to successfully prey on eared moth species. Similar to Shiel et al. (1991), this study found that Diptera played an important part of the diet of this species in all roosts. This ability to feed on Diptera species was also confirmed by Roswag et al. (2018). The flexibility in prey selection was further illustrated at one roost location (Kilmore), where individuals of the roost fed more on Diptera than Lepidoptera. In highly protected areas like Glengarriff (the roost sampled is located within the Glengarriff Nature Reserve), the diet included a higher proportion of Coleoptera (10.5%), Aranea (5.1%), and Lithobiomorpha (8.2%), indicating that habitat quality might influence prey composition. This flexibility was also reflected by the diet distinctiveness at the species level across roosts. Our findings indicate that the Brown long-eared bat demonstrates flexibility in its foraging behaviour and can adapt its diet depending on prey availability as already suggested (Rostovskaya et al., 2000). Although often considered as a Lepidoptera specialist (Dietz and Kiefer, 2016; Wilson and Mittermeier, 2019; Ancillotto and Russo, 2023), our findings support a broader foraging strategy, positioning the Brown long-eared bat as a generalist with a preference for Lepidoptera.

Prey richness varied significantly between faecal samples, averaging 6.9 species per sample. This is notably lower than the richness observed by Andriollo et al. (2019), who reported between 21.8 and 26.9 species per sample in Switzerland. However, Shiel et al. (1998), found lower prey diversity in *Nyctalus leisleri* faecal samples in Ireland compared to those from Germany. These differences in prey richness may reflect unique aspects of the Irish landscape and habitat composition.

### 4.2 Soprano pipistrelle’s diet

The Soprano pipistrelle exhibited a similarly broad diet, feeding on prey from eleven arthropod orders, with 350 taxa identified at the species level. In contrast to the Brown long-eared bat, Soprano pipistrelles showed a strong preference for Diptera species, accounting for 60% of the species identified. Food web analyses further emphasised this reliance, as Diptera interactions comprised over 83% of total prey interactions. The prevalent Diptera families included Psychodidae, Chironomidae, Limoniidae, Tipulidae, and Anisopodidae. This aligns with previous findings of Puig-Montserrat et al. (2020) who also reported a high-occurrence of these families in the Soprano pipistrelle diet in Catalogna, Spain. However, unlike this study that identified Drosophilidae as a dominant component of the diet, our study revealed infrequent interactions with this family in Irish Soprano pipistrelles, detecting it at only two sites. The Soprano pipistrelle’s diet revealed notable flexibility within Diptera, with prey covering a wide range of families and species, suggesting adaptability in foraging behavior depending on prey availability. Additionally, while the majority of the prey had small forewing length, such as Psychodidae (2–6 mm) and Chironomidae (1–10 mm), larger prey like Tipulidae were also consumed, including species like *Tipula maxima* (22–30 mm), *Tipula paludosa* (13–23 mm), and *Tipula oleracea* (18–28 mm) (Brock, 2020). This wide range of prey sizes demonstrates that the Soprano pipistrelle is not limited to a narrow prey size range.

Similar to the Brown long-eared bat, the mean prey richness per faecal sample was relatively low (seven species), though variation was observed across roosts and sampling years. This variation may reflect fluctuations in local prey availability, seasonal shifts in prey populations, or environmental factors influencing foraging behavior.

Due to the limited number of studies focusing on the diet of the Soprano pipistrelle, our findings provide important new insights into its feeding ecology. However, it is important to acknowledge that our study may have underestimated the full diversity of its diet. The small number of maternity roosts included in the study, combined with potential limitations in sampling coverage, could have resulted in the underrepresentation of rare or less frequently consumed prey. Nevertheless, the roosts sampled were situated in areas representing a good cross-section of Irish landscapes, which likely helped reduce this bias.

### 4.3 Influence of land cover, year, and time

The richness of the bat species’ diet was influenced by the species itself, the roost location, and the year of sampling. However, our analyses did not clearly identify which of these factors had the strongest influence on dietary richness, likely due to the limited number of roosts sampled. The variations observed in richness over time might reflect fluctuating prey availability (Clare et al., 2011 ; Clare et al., 2014) though no clear temporal pattern emerged for either Lepidoptera or Diptera species. A notable observation was made at the Killmore roost, where individuals consumed more Diptera species than at other roosts. This may be attributed to a higher local abundance of Diptera or a lower availability of Lepidoptera, causing a dietary shift. While the land cover around Kilmore is dominated by improved grassland (62.5%), similar trends were not observed at other roosts with comparable land cover percentages. Interestingly, bats at this site consumed more Muscidae and Scathophagidae species, suggesting a site-specific influence on diet composition. Improved grasslands are known to support lower arthropod diversity (Arnott et al., 2022), which may contribute to this shift; however, the specific factors behind the increased Diptera consumption at Killmore remain unclear. Importantly, across all sampled roosts and years, the prey composition detected in faecal samples was lower in prey richness compared to studies conducted in more structurally diverse or heterogeneous landscapes (Andriollo et al., 2019).

This suggests that bats foraging in pastureland-dominated habitats may encounter reduced prey diversity, potentially constraining their foraging behaviour. However, Ireland’s comparatively low insect diversity relative to other European regions (McCarthy, 1986; Nelson et al., 2011; Kelly-Quinn and Regan, 2012; Harrison, 2014; Feeley et al., 2020) may also have contributed to these findings. These findings emphasised the importance of considering landscape structure and arthropod biodiversity when assessing bat dietary ecology and support the idea that habitat complexity can enhance prey availability and trophic niche width.

Roost location also significantly impacted diet composition for both bat species. However, unlike the findings of Tournayre et al. (2021), who reported a strong correlation between landscape dissimilarity and diet composition in Greater horseshoe bats (*Rhinolophus ferrumequinum*), our results showed only a small positive correlation between these factors. The difference in findings may be attributed to differences in the species studied and their foraging habits.

Hierarchical clustering analysis indicated that the temporal factor (year of sampling) had a notable effect on diet composition. Nonetheless, the relative influence of land cover and year differed among maternity roosts. In some cases, interannual variations had a stronger influence on diet composition than spatial location, while in others, spatial factors were more influential. This suggests that both temporal and spatial factors contribute to shaping bat diets, likely due to their both influence on prey availability (Uhler et al., 2021; Saha et al., 2023). Nevertheless, with our current dataset, it remains difficult to discern which of these is the dominant driver. A strong impact of temporal dynamics has previously been reported for the Brown long-eared bat (Razgour et al., 2011; Andriollo et al., 2019). However, our results indicate that at several maternity roosts, the effect of year on diet composition appeared weaker than land cover, suggesting that local habitat characteristics may have stronger influence on diet patterns than intra-annual variations in specific contexts. The key drivers, however, remain unclear and may reflect complex interactions between local prey availability, habitat structure, and bat foraging behaviour.

### 4.4 Limitations

Despite the strengths of the methodological approach, several limitations must be considered. In this study, a presence/ absence approach was used instead of a relative abundance approach. Presence/absence data give more weight to low-frequency prey taxa (Taberlet et al., 2018; Deagle et al., 2019) while the read abundance is influenced by biases introduced during DNA extraction, PCR amplification, and sequencing biases (Elbrecht and Leese, 2015; Pawluczyk et al., 2015; Taberlet et al., 2018; Yang et al., 2021; Martoni et al., 2022).

Therefore, the presence/absence approach was considered more appropriate for this study.

We implemented stringent bioinformatic filtering, excluding MOTUs representing less than 1% of sequences per faecal sample to avoid contamination or secondary predation. While this may have excluded some rare prey items (Mata et al., 2021; Alberdi et al., 2018), we believe it provided a reliable picture of the main dietary components for both bat species.

A key limitation lies in taxonomic assignment, which depends on the completeness of reference databases. Although we used two CO1 gene regions to improve taxonomic resolution, geographic scale can induce intraspecific genetic variation in COI sequences (Bergsten et al., 2012), which may have resulted in some MOTUs being unassigned or misassigned to closely related species. This highlights the need for an Ireland-specific arthropod DNA database, as current global databases may not fully represent the local biodiversity. Additionally, only species documented as present in Ireland were included in the final analysis. As a result, undocumented prey species may have been excluded from the analysis.

Additionally, sample coverage exceeded 91% overall, but some roosts and sampling events fell below 90%. This lower coverage could have led to an underestimation of dietary diversity at certain sites or times, suggesting that a small fraction of the diet may not have been detected in our analysis.

### 4.5 Conservation implications

Our findings underline the importance of conserving diverse insect populations, particularly Lepidoptera and Diptera, which are crucial to the diets of both the Brown long-eared bat and Soprano pipistrelle. Agricultural intensification, habitat degradation, and pesticide use pose serious threats to these prey groups (Wagner, 2020 ; Rumohr et al., 2023), potentially reducing prey availability and impacting bat populations.

Although both bat species demonstrated flexibility in their diets, the Soprano pipistrelle’s strong reliance on Diptera may heighten its vulnerability to ongoing declines in arthropod populations (Rumohr et al., 2023). This highlights the need for targeted conservation strategies that safeguard both prey diversity and habitat quality.

Bats also provide important ecosystem services, such as pest control, by regulating insect populations, including agricultural pests (Puig-Montserrat et al., 2020 ; Tiede et al., 2020 ; Charbonnier et al., 2021 ; Montauban et al, 2021 ; Da Silva et al., 2023; Hunnick et al., 2022). Understanding the role of Irish bats in these services could strengthen conservation efforts by highlighting their ecological and economic importance. Future research should focus on quantifying these ecosystem services and developing a comprehensive Irish arthropod DNA database to improve metabarcoding accuracy. Additionally, studies on the effects of climate change and habitat alteration on bat diets and prey availability will be crucial for informing long-term conservation strategies.

## 5. Conclusion

Using metabarcoding and next-generation sequencing on faecal samples collected over three years, this study offers new insights into the diet of the Brown long-eared bat and the Soprano pipistrelle in pastureland-dominated landscapes. It confirms the previously suggested foraging flexibility of the Brown long-eared bat and the high reliance of the Soprano pipistrelle on Diptera prey, in a habitat where pasture lands predominate. Interestingly, prey richness in faecal samples across all roosts and both bat species was lower than reported in comparable studies from different or more structurally complex landscapes. This study offers a strong foundation for understanding the feeding ecology of bats in pastureland dominated landscapes and contributes to a broader understanding of how landscape context influences bat foraging ecology. Continued research is essential for understanding bat-prey interactions better and for developing conservation strategies that protect both bats and their prey, as well as the important ecosystem services they provide, in a changing world.

## Acknowledgements

This research was supported by the Irish Research Council of Ireland Postgraduate Scholarship (grant no. GOIPG/2020/1517) awarded to G.H. and E.C.T. and National Parks and Wildlife Service (NPWS) of Ireland funding awarded to E.C.T.

We would like to thank James Nolan, Sarah Redmond, and Dr. Sébastien Riquier for their invaluable assistance during fieldwork, and Dr. Aidan O’Hanlon (National Museum of Ireland) for his helpful support in validating the presence of the Irish arthropods identified by our metabarcoding.

## Author contributions

Gwenaëlle Hurpy, Niamh Roche, and Emma C. Teeling designed the project. GH and Tina Aughney organised and collected bat faecal samples. GH and Ilze Skujina conducted the laboratory work. Statistical analysis and figures were generated by GH with input of ECT. GH drafted the manuscript with input from ECT. All authors critically revised and edited the manuscript. All authors gave their final approval of the manuscript to be published.

## Data accessibility and benefit-sharing

Raw sequence and metadata are available on DataDryad (DOI: https://doi.org/10.5061/dryad.qbzkh18vt)

