## Supporting Information for "Metabarcoding reveals the effect of land cover and temporal variations on the diet of Brown long-eared and Soprano pipistrelle bats within a unique European habitat, the pasture dominated landscapes of Ireland"

***Table of Contents***

| | |
|---|---|
| ***Page 2*** | ***Supporting Table S1*** |
| ***Page 3*** | ***Supporting Table S2*** |
| ***Page 4*** | ***Supporting Figure S1*** |
| ***Page 15*** | ***Supporting Figure S2*** |
| ***Page 16*** | ***Supporting Figure S3*** |
| ***Page 19*** | ***Supporting Table S3*** |
| ***Page 20*** | ***Supporting Figure S4*** |

***Supporting Table S1: Number of faecal samples collected at each bat maternity roost over the three years of sampling (2021-2023).*** *$1^{st}$ sampling period : Gestation ; $2^{nd}$ sampling period : Lactation ; 3rd sampling period : Post-lactation. Numbers of faecal samples included in the analyses, after that bat species was confirmed, are indicated between parentheses. The sub-total lines show the total of number of faecal samples collected per bat species. The total line presents the combined total for the two bat species. The numbers indicated in the "roost column" correspond to the numbers assigned to each roost in Fig. 1.*

| | | **2021 sampling collections** | | | **2022 sampling collections** | | | **2023 sampling collections** | | | **TOTAL** |
|---|---|---|---|---|---|---|---|---|---|---|---|
| **Bat species** | **Roost** | **$1^{st}$** | **$2^{nd}$** | **$3^{rd}$** | **$1^{st}$** | **$2^{nd}$** | **$3^{rd}$** | **$1^{st}$** | **$2^{nd}$** | **$3^{rd}$** | |
| **Soprano pipistrelle *Pipistrellus pygmaeus*** | Glendalough (1) | 16(16) | 135(20) | 77(20) | 25(20) | 42(20) | 91(20) | 23(20) | 45(20) | 36(20) | **490(176)** |
| | Kilafin (2) | / | / | / | 7(7) | 13(13) | 11(10) | 5(1) | 20(19) | 23(20) | **79(70)** |
| | Birr (3) | 30(0) | 41(12) | 36(15) | 11(7) | 26(13) | 15(10) | 45(20) | 23(20) | 12(12) | **239(109)** |
| | Dromore wood (4) | / | / | / | 54(20) | 72(20) | 29(20) | 30(20) | 40(20) | 28(20) | **253(120)** |
| | **Sub-total** | **46(16)** | **176(32)** | **113(35)** | **97(54)** | **153(66)** | **146(61)** | **103(61)** | **128(79)** | **99(72)** | **1061(475)** |
| **Brown long-eared bat *Plecotus auritus*** | Ennisnag (5) | 31(17) | 10(2) | 6(5) | 43(16) | 36(20) | 20(6) | 17(14) | 33(12) | 27(15) | **223(107)** |
| | Glengarriff (6) | 39(18) | 47(16) | 52(12) | 19(18) | 11(11) | 22(20) | 50(20) | 38(20) | 63(18) | **341(153)** |
| | Inagh (7) | 197 (20) | 103(20) | 97(20) | 11(11) | 119(20) | 55(20) | 56(20) | 46(20) | 48(20) | **732(171)** |
| | Kinvarra (8) | 78(20) | 108(20) | 168(20) | 68(20) | 171(20) | 76(20) | 79(20) | 33(20) | 71(20) | **852(180)** |
| | Letterfrack (9) | 11(10) | 19(18) | 182(19) | 20(20) | 52(20) | 64(20) | 54(20) | 97(20) | 45(20) | **544(167)** |
| | Carra James (10) | 43(20) | 46(20) | 30(20) | 26(20) | 30(20) | 30(20) | 33(20) | 27(20) | 23(20) | **288(180)** |
| | Milltown (11) | 35(19) | 26(0) | 48(17) | 26(20) | 30(20) | 30(20) | 30(20) | 34(20) | 38(20) | **297(156)** |
| | Killmore (12) | 11(11) | 96(16) | 17(17) | 21(17) | 54(20) | 22(14) | 12(12) | 24(16) | 32(20) | **289(143)** |
| | **Sub-total** | **445 (135)** | **455 (112)** | **600 (130)** | **234 (142)** | **503 (151)** | **319 (140)** | **331 (146)** | **332 (148)** | **347 (153)** | **3566 (1257)** |
| | **TOTAL** | **491 (151)** | **631 (145)** | **713 (165)** | **331 (196)** | **656 (217)** | **465 (201)** | **434 (207)** | **460 (228)** | **446 (225)** | **4627 (1732)** |

***Supporting Table S2: NucleoSpin Plant kit extraction protocol (adapted from Zarzoso-Lacoste et al., 2018).***

| **NucleoSpin Plant kit extraction protocol<br>(adapted from Zarzoso-Lacoste *et al*., 2018)** |
|---|
| **1. Sample preparation** |
| Add carefully 0.01 gr of pooled faeces into 2mL tube |
| **2. Cell lysing Buffer PL2 and PL3** |
| Add 400 µL Buffer PL2 and 10 µL RNase to each tube strips containing faeces sample and close using new cap strips. Mix by vigorous shaking for 15–30 s. Spin for 30 s at 11 000 rpm to collect any sample from the Cap Strips. |
| Incubate samples at 60°C overnight, on dry bath. |
| Carefully add 100 µL Buffer PL3 to each sample and close the tubes, mix thoroughly, and incubate for 15 min on ice to precipitate sodium dodecyl sulfate completely. |
| **3. Filtrate lysate** |
| Place NucleoSpin filter (violet ring) into new collection tube 2mL |
| Load all lysate and centrifuge for 2min at 11 000 g. |
| Collect filtered lysate into a new 1.5 Eppendorf tube and discard filter |
| **4. Adjust DNA binding condition** |
| Predispose 450 µL Buffer PC in 1.5mL Eppendorf |
| Add 400 µL clear lysate and mix thoroughly by pipetting five times |
| **5. Bind DNA** |
| Label green column and place it on new column |
| Transfer 700 µL of lysate to column and centrifuge for 2 min at 14 000 rpm and discard flow through |
| **6. Wash and dry silica membrane** |
| Add 400 µL PW1 Buffer and centrifuge for 1 min at 14 000 rpm |
| Add 700 µL PW2 Buffer and centrifuge for 1 min at 14 000 rpm |
| Add 700 µL PW2 Buffer and centrifuge for 1 min at 14 000 rpm |
| Centrifuge at 14 000 rpm for 10 min |
| **7. Elute DNA** |
| Place column in labelled 1.5 Eppendorf |
| Dispose PE Buffer 75 µL into column (pre-heated at 70°C) and incubate at 70°C for 2 min |
| Centrifuge 2 min at 14 000 rpm |
| Dispose PE Buffer 75 µL into column (pre-heated at 70°C) and incubate at 70°C for 2 min |
| Centrifuge 2 min at 14 000 rpm |
| **Dispose in - 20°C freezer of use for PCR immediately** |

***Supporting Figure S1. Sample completeness curves for each maternity roost by year and for each sampling period.*** *Curves were created with iNEXT and ggiNEXT packages, with q = 0.  Rarefaction curve (solid line) represents the expected sample completeness as the sample size increases. Extrapolation curve (dashed line) predicts the potential sample completeness beyond the observed sample size. A - H : Brown long-eared bat roosts; I - L : Soprano pipistrelle roosts..*

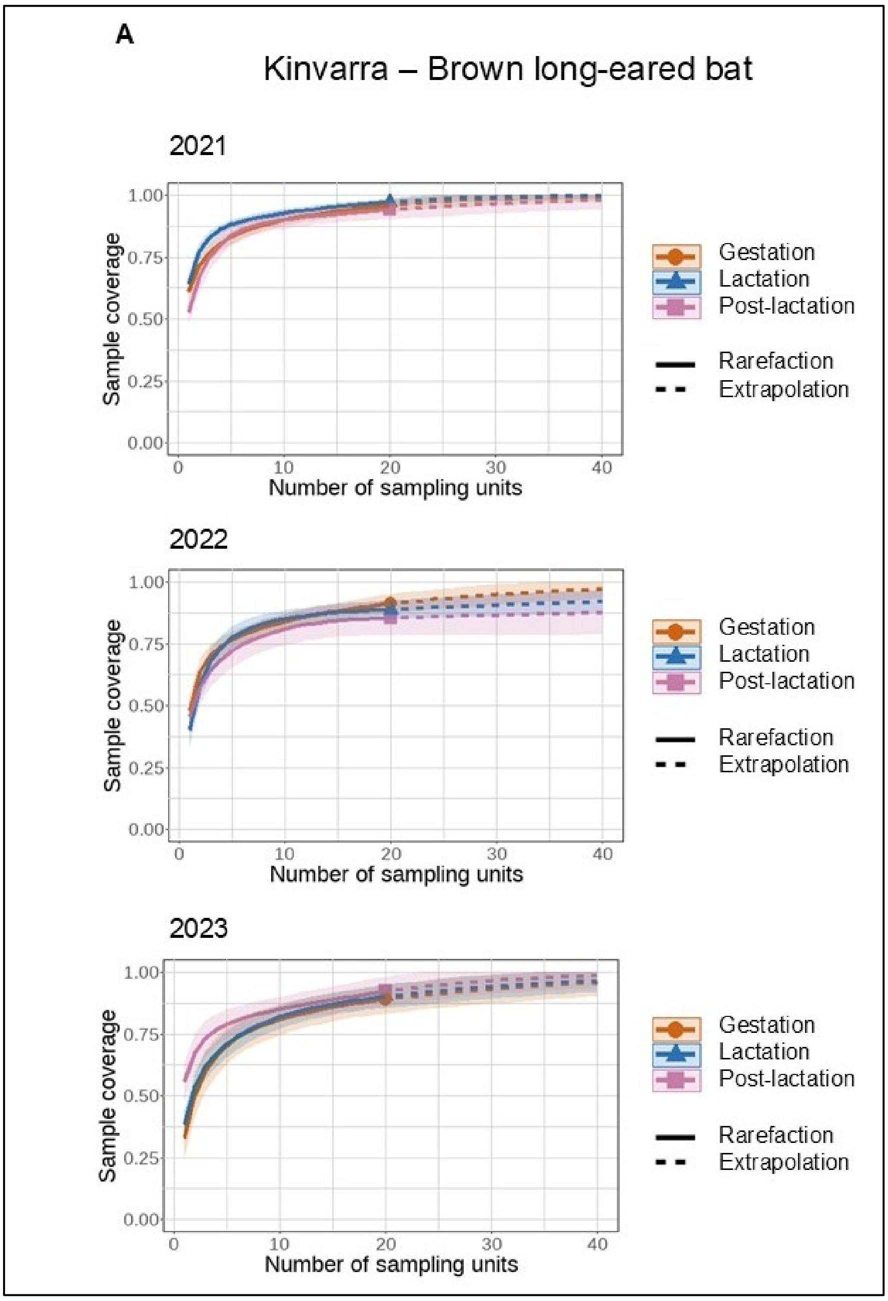

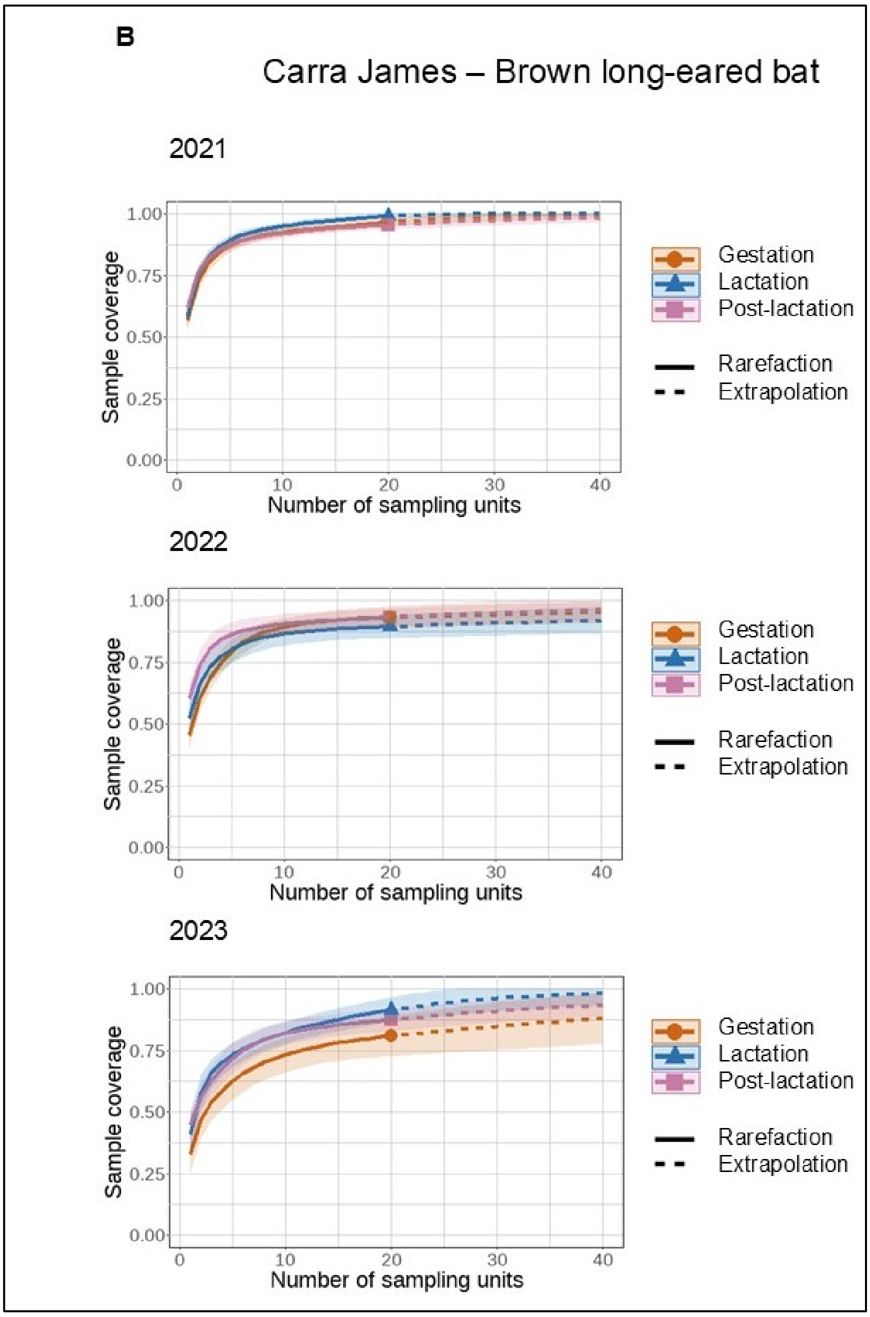
B
Carra James – Brown long-eared bat
2021
2022
2023
Sample coverage
Number of sampling units
Gestation
Lactation
Post-lactation
Rarefaction
Extrapolation

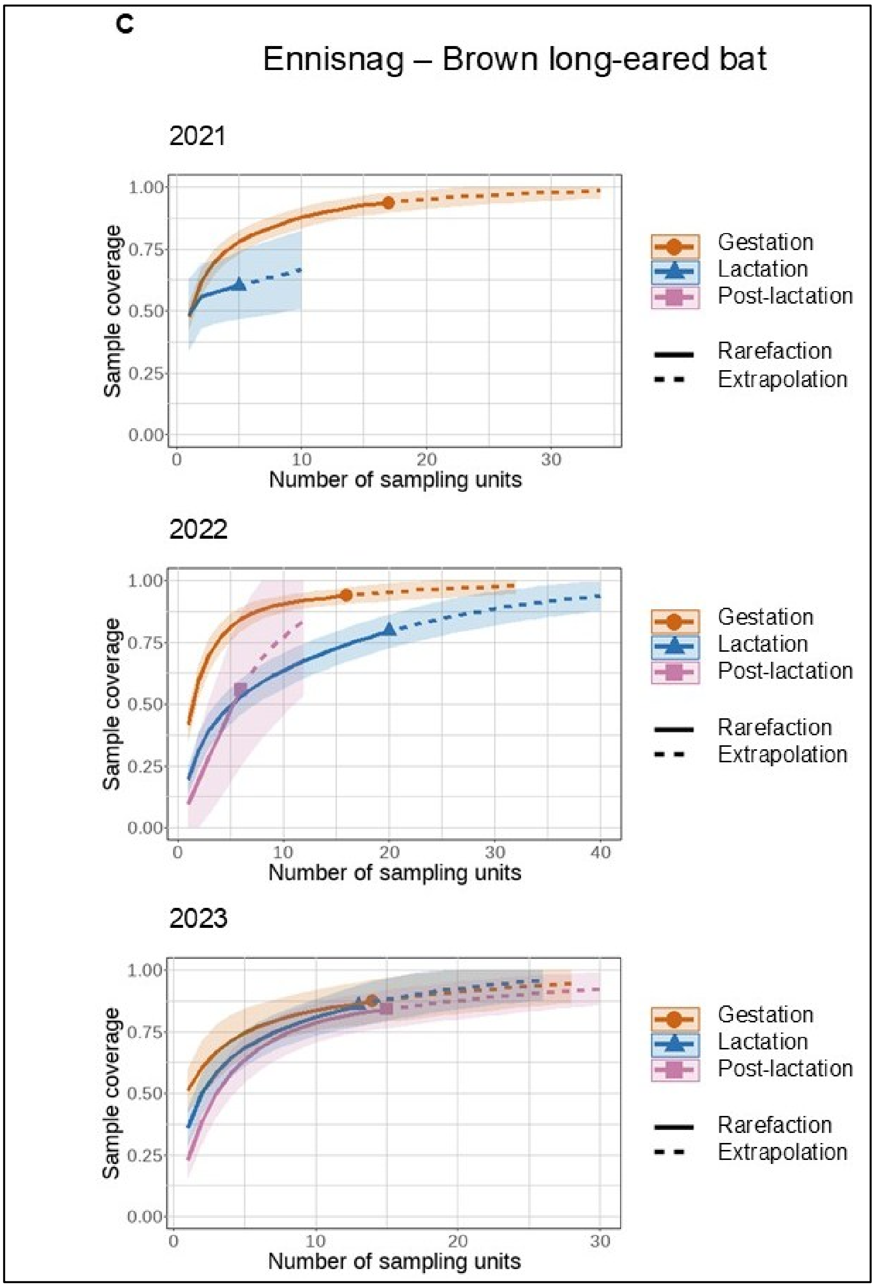

C
Ennisnag – Brown long-eared bat
2021
2022
2023
Sample coverage
Number of sampling units
1.00
0.75
0.50
0.25
0.00
0
10
20
30
40
Gestation
Lactation
Post-lactation
Rarefaction
Extrapolation

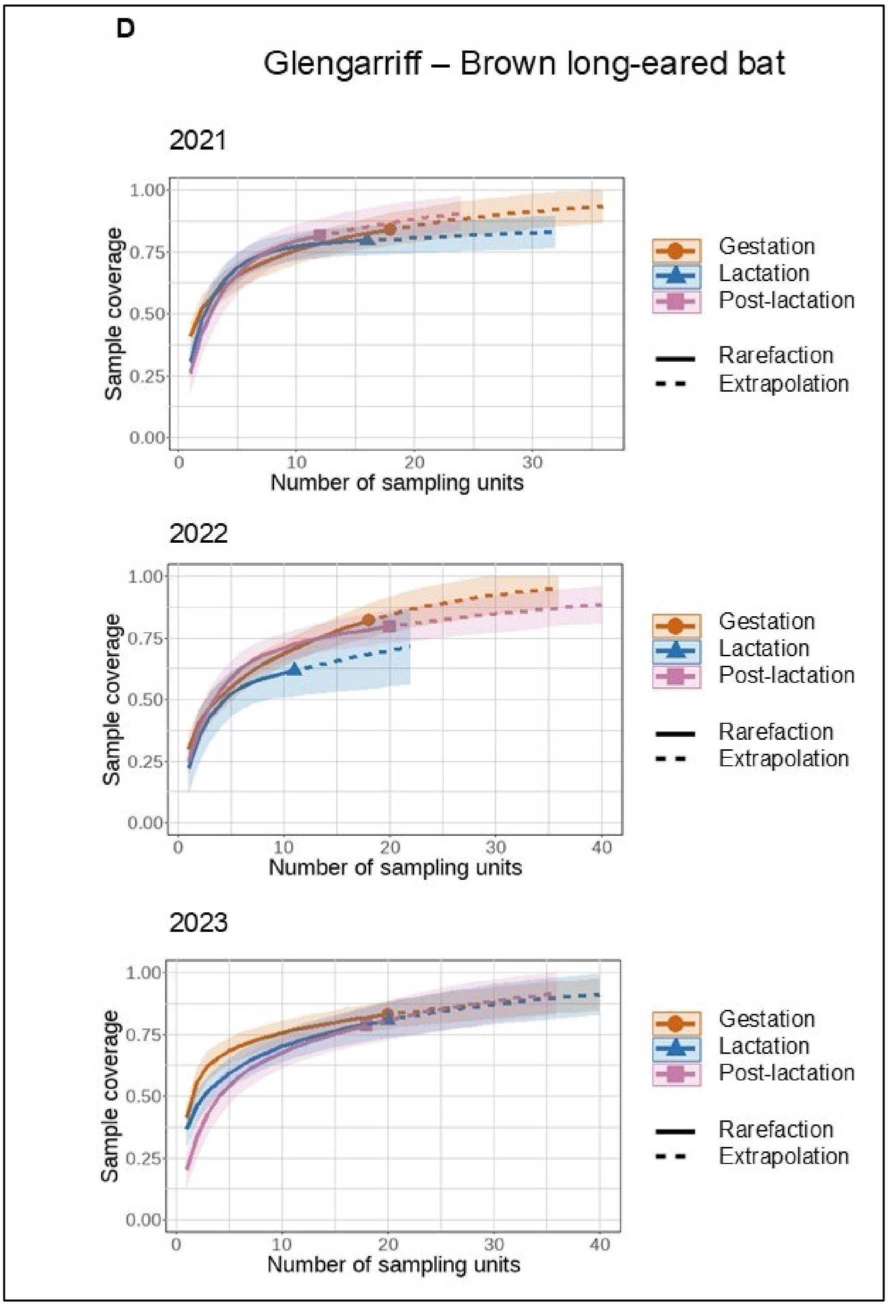

D
Glengarriff – Brown long-eared bat
2021
Sample coverage
Number of sampling units
Gestation
Lactation
Post-lactation
Rarefaction
Extrapolation
2022
Sample coverage
Number of sampling units
Gestation
Lactation
Post-lactation
Rarefaction
Extrapolation
2023
Sample coverage
Number of sampling units
Gestation
Lactation
Post-lactation
Rarefaction
Extrapolation

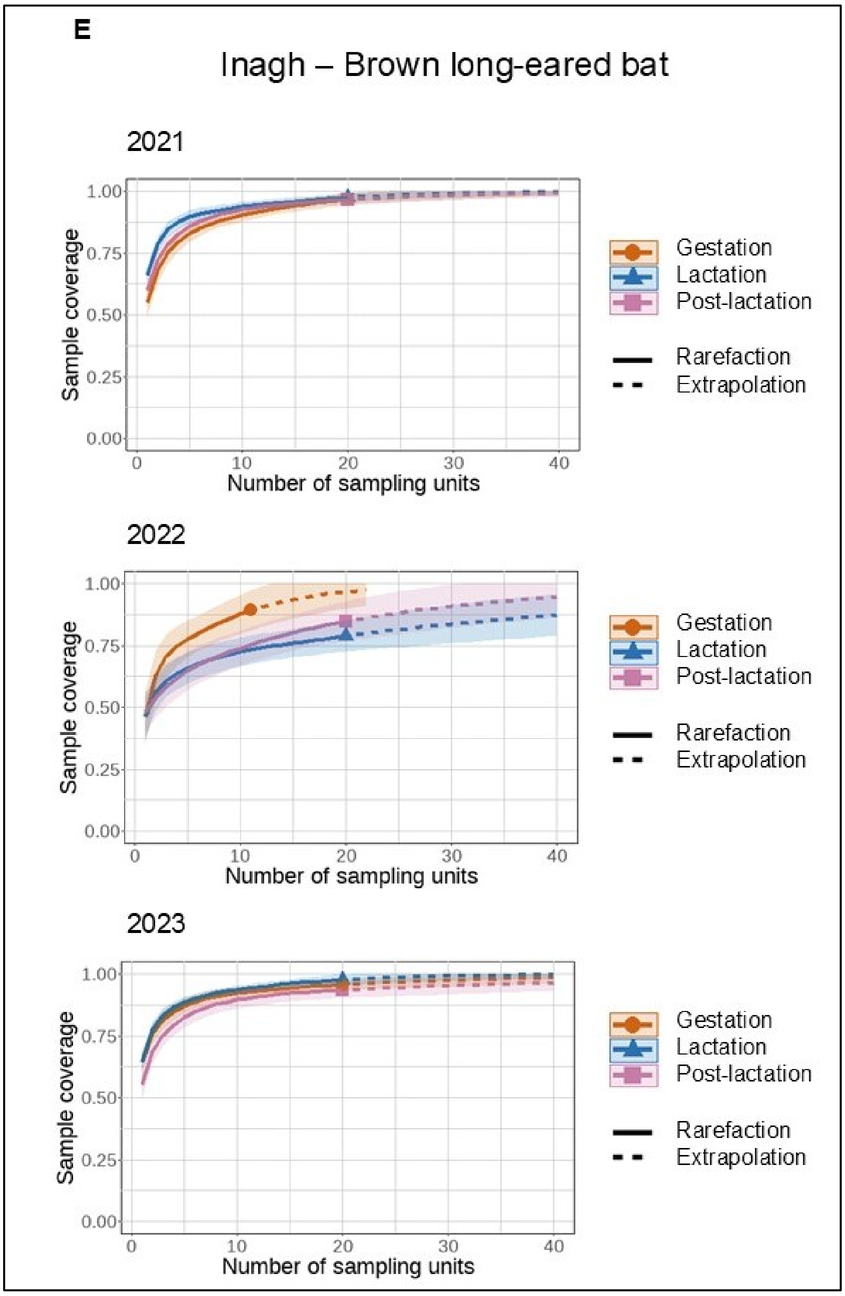
E
Inagh – Brown long-eared bat
2021
2022
2023
Sample coverage
1.00
0.75
0.50
0.25
0.00
0
10
20
30
40
Number of sampling units
Gestation
Lactation
Post-lactation
Rarefaction
Extrapolation

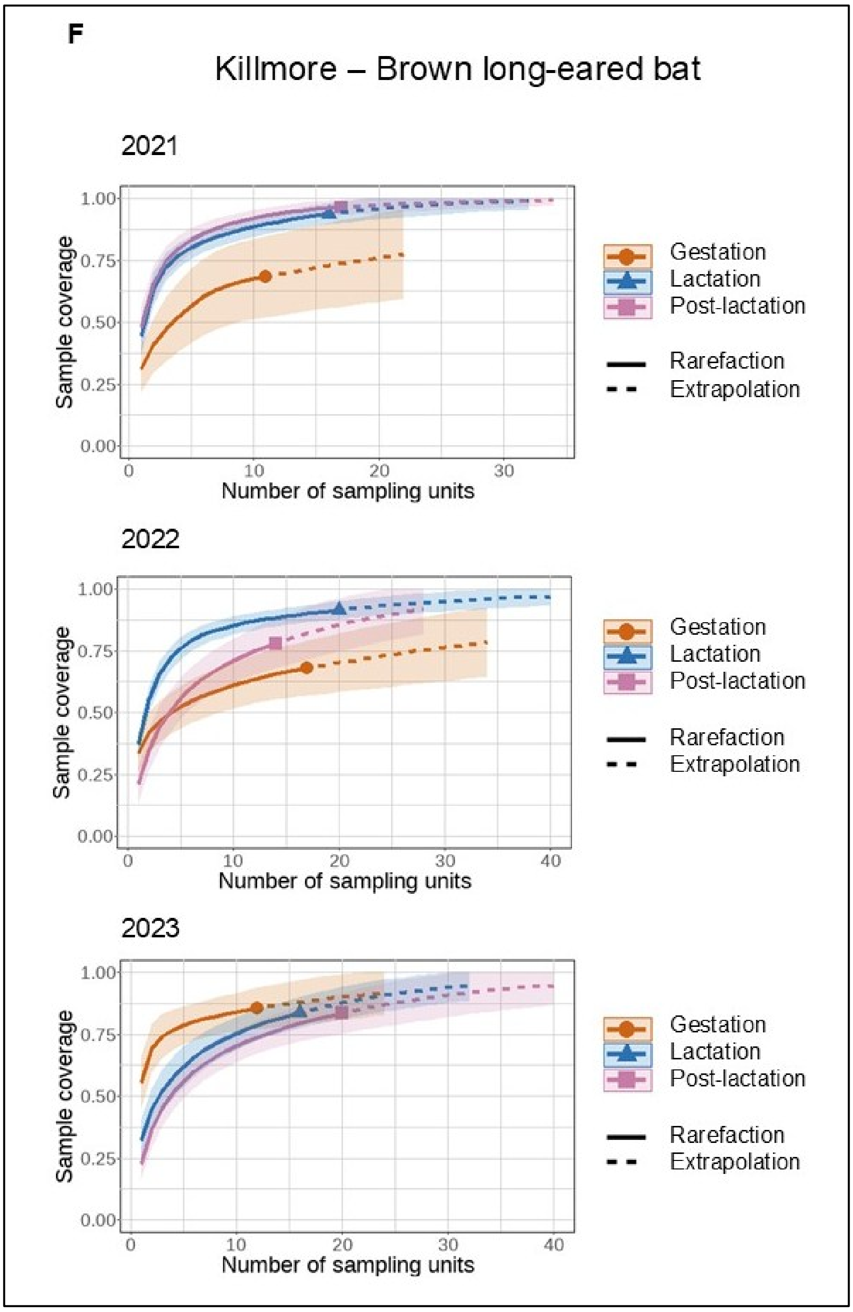
F
Killmore – Brown long-eared bat
2021
2022
2023
Sample coverage
Number of sampling units
Gestation
Lactation
Post-lactation
Rarefaction
Extrapolation

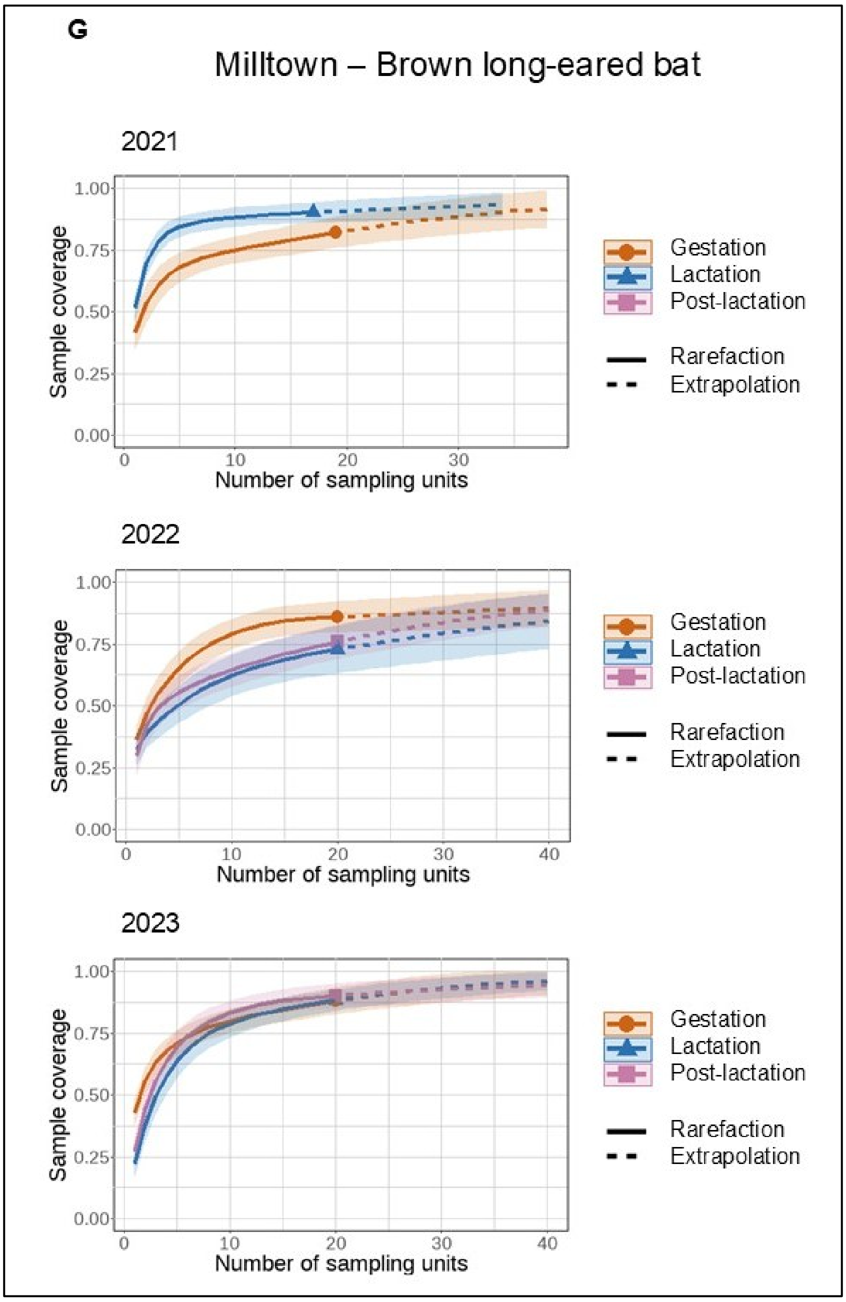
G
Milltown – Brown long-eared bat
2021
2022
2023
Sample coverage
Number of sampling units
1.00
0.75
0.50
0.25
0.00
0
10
20
30
40
Gestation
Lactation
Post-lactation
Rarefaction
Extrapolation

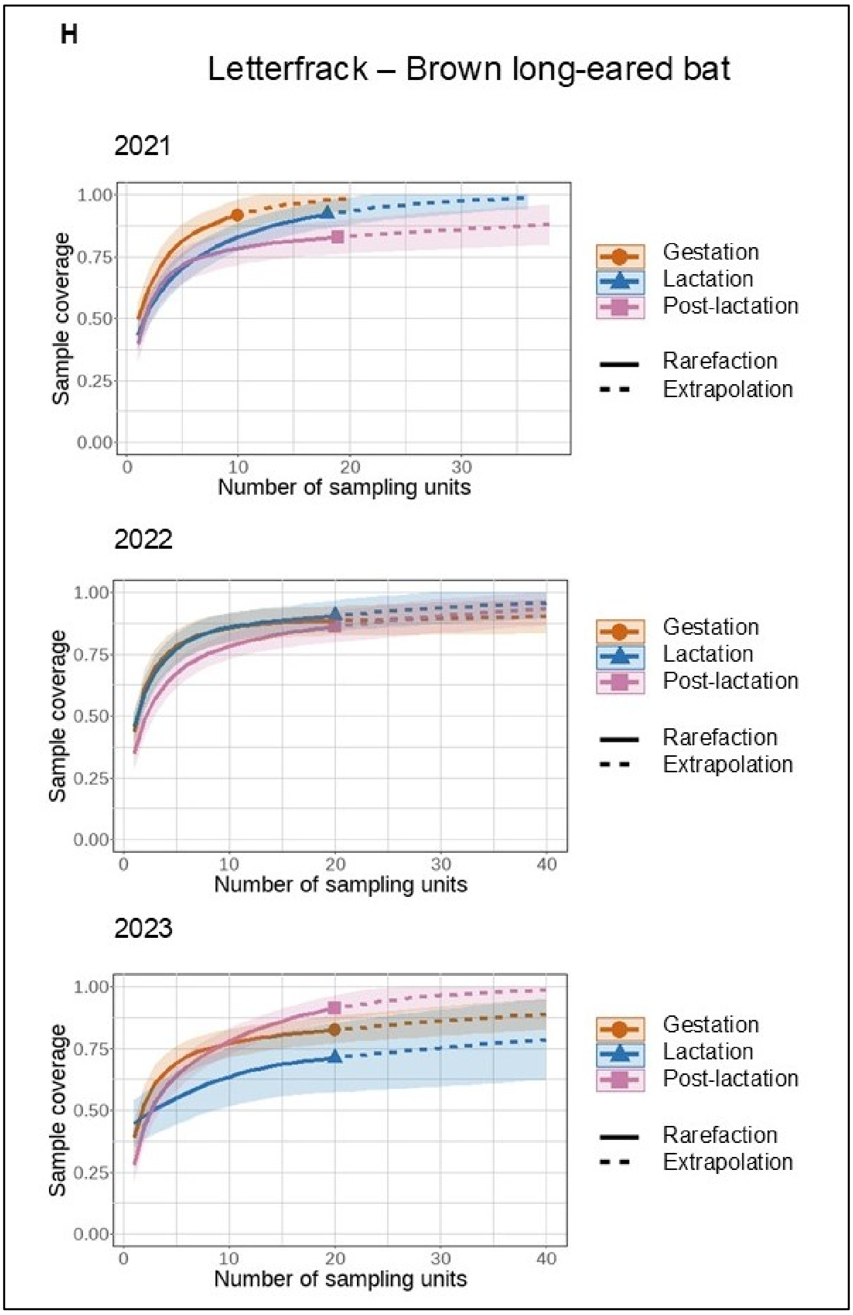
H
Letterfrack – Brown long-eared bat
2021
2022
2023
Sample coverage
Number of sampling units
Gestation
Lactation
Post-lactation
Rarefaction
Extrapolation

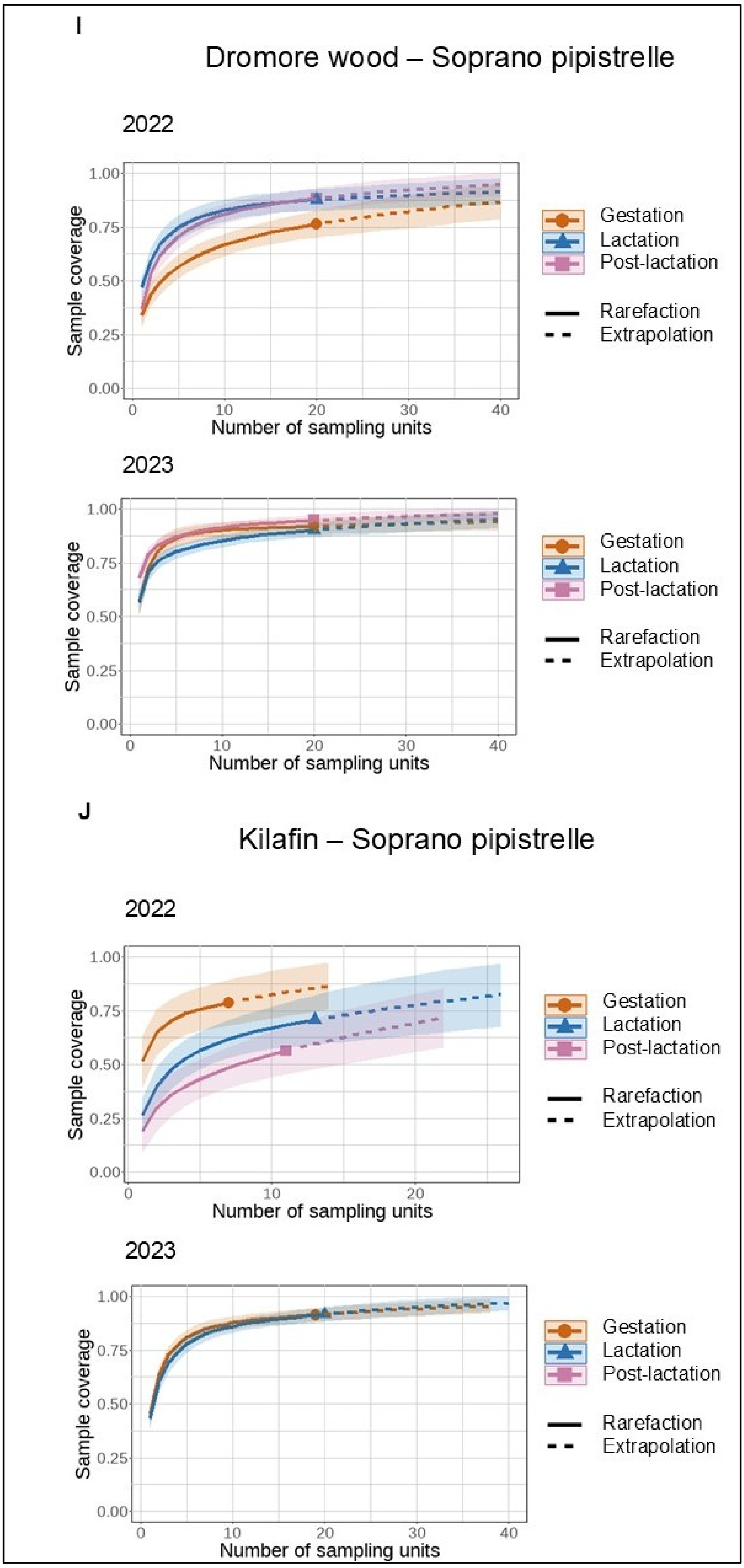
I
Dromore wood – Soprano pipistrelle
2022
Sample coverage
Number of sampling units
Gestation
Lactation
Post-lactation
Rarefaction
Extrapolation
2023
Sample coverage
Number of sampling units
Gestation
Lactation
Post-lactation
Rarefaction
Extrapolation
J
Kilafin – Soprano pipistrelle
2022
Sample coverage
Number of sampling units
Gestation
Lactation
Post-lactation
Rarefaction
Extrapolation
2023
Sample coverage
Number of sampling units
Gestation
Lactation
Post-lactation
Rarefaction
Extrapolation

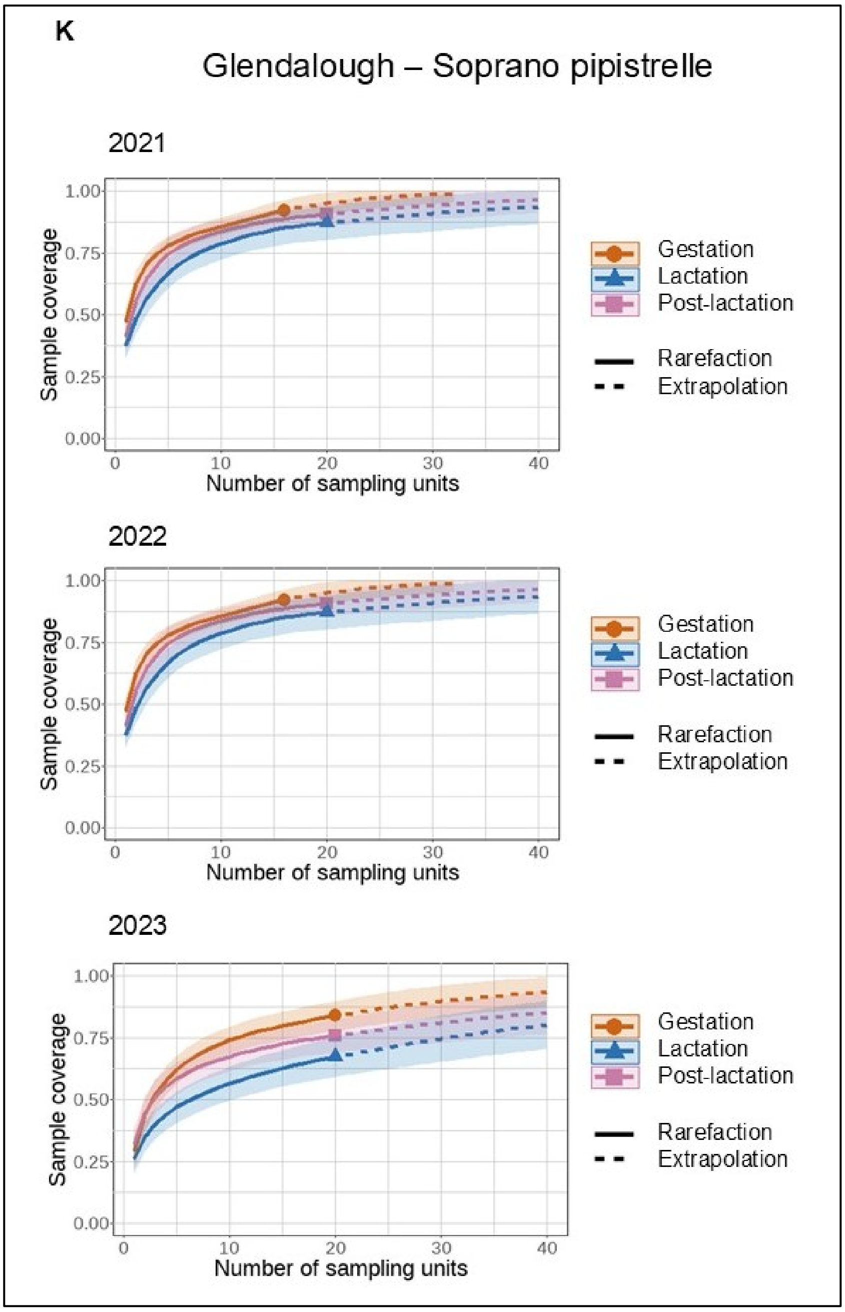
K
Glendalough – Soprano pipistrelle
2021
Sample coverage
Number of sampling units
Gestation
Lactation
Post-lactation
Rarefaction
Extrapolation
2022
Sample coverage
Number of sampling units
Gestation
Lactation
Post-lactation
Rarefaction
Extrapolation
2023
Sample coverage
Number of sampling units
Gestation
Lactation
Post-lactation
Rarefaction
Extrapolation

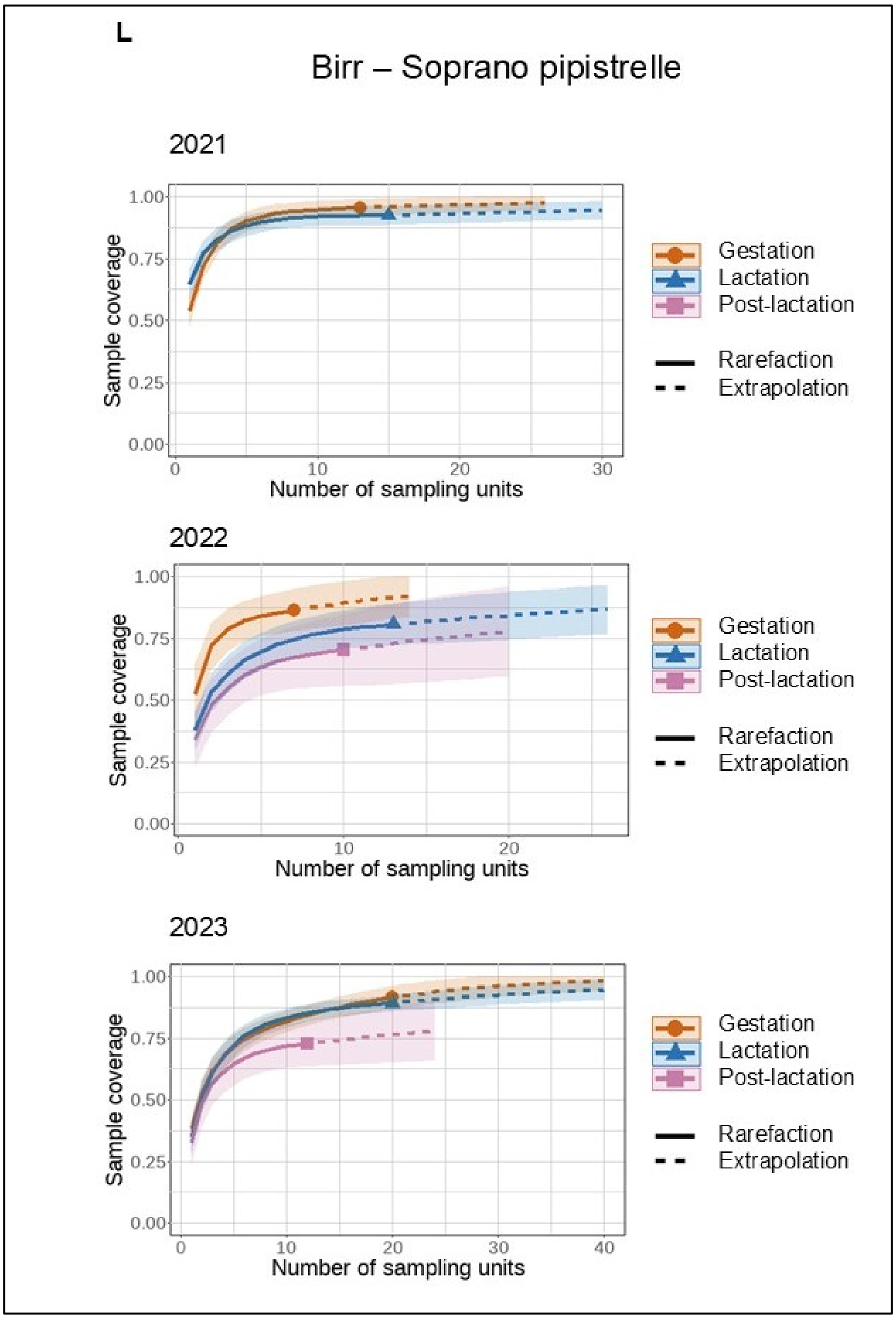
L
Birr – Soprano pipistrelle
2021
Sample coverage
Number of sampling units
Gestation
Lactation
Post-lactation
Rarefaction
Extrapolation
2022
Sample coverage
Number of sampling units
Gestation
Lactation
Post-lactation
Rarefaction
Extrapolation
2023
Sample coverage
Number of sampling units
Gestation
Lactation
Post-lactation
Rarefaction
Extrapolation

***Supporting Figure S2***

***Mean richness of main arthropod orders (Lepidoptera and Diptera) in the diet of Brown long-eared bat (A) and Soprano pipistrelle (B) across all sampling periods****. Horizontal lines at each point represent standard deviation values*

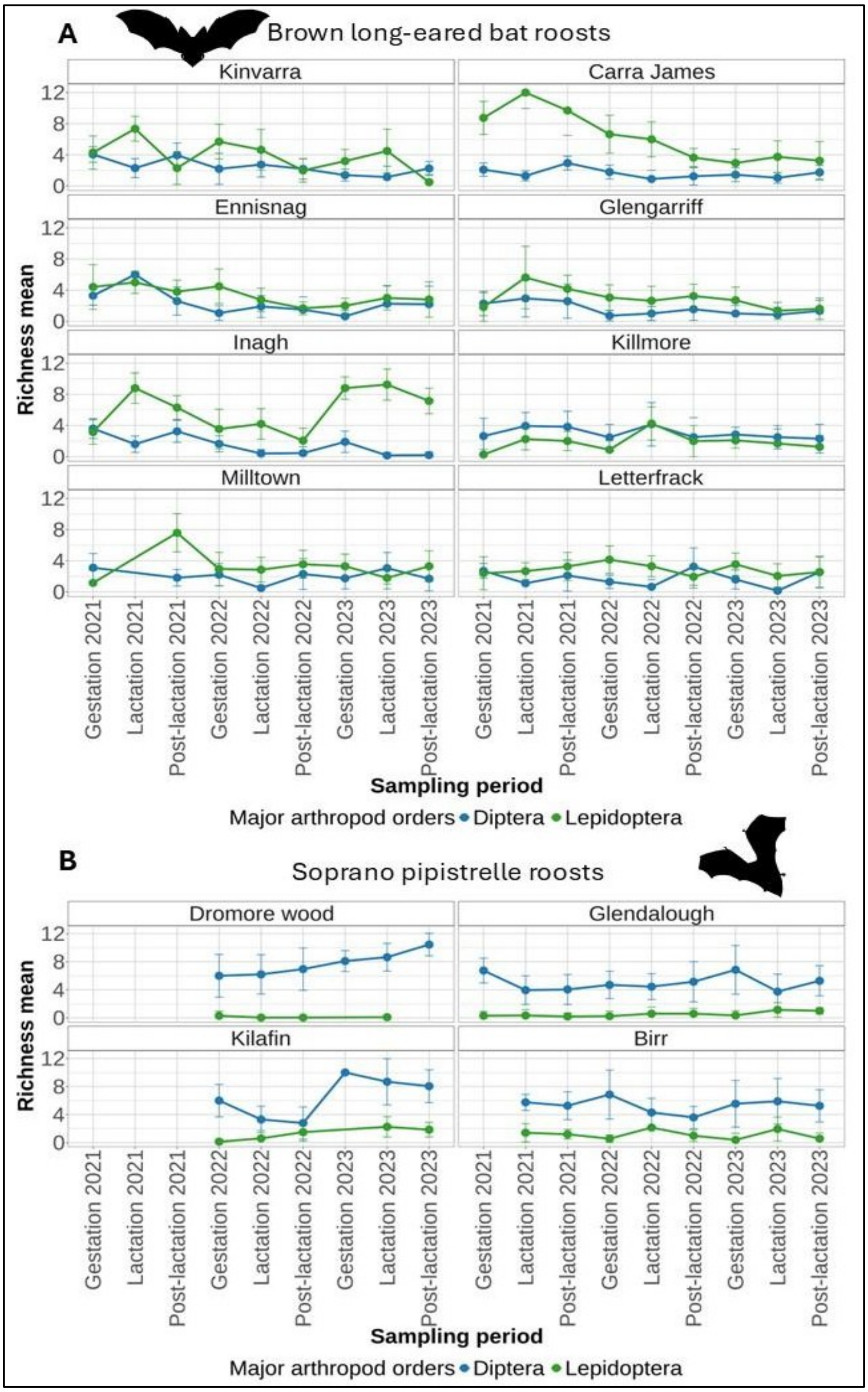

**Supplementary Figure S3**

***Web plots showing interactions between the Brown long-eared bat (S.3A) and Soprano Pipistrelle (S.3B) and prey of Lepidoptera (a) and Diptera (b) families across maternity roosts****. Percentages show the percentage of each order in diets. The web plot was created using the ‘bipartiteD3’ function from the bipartiteD3 package*.

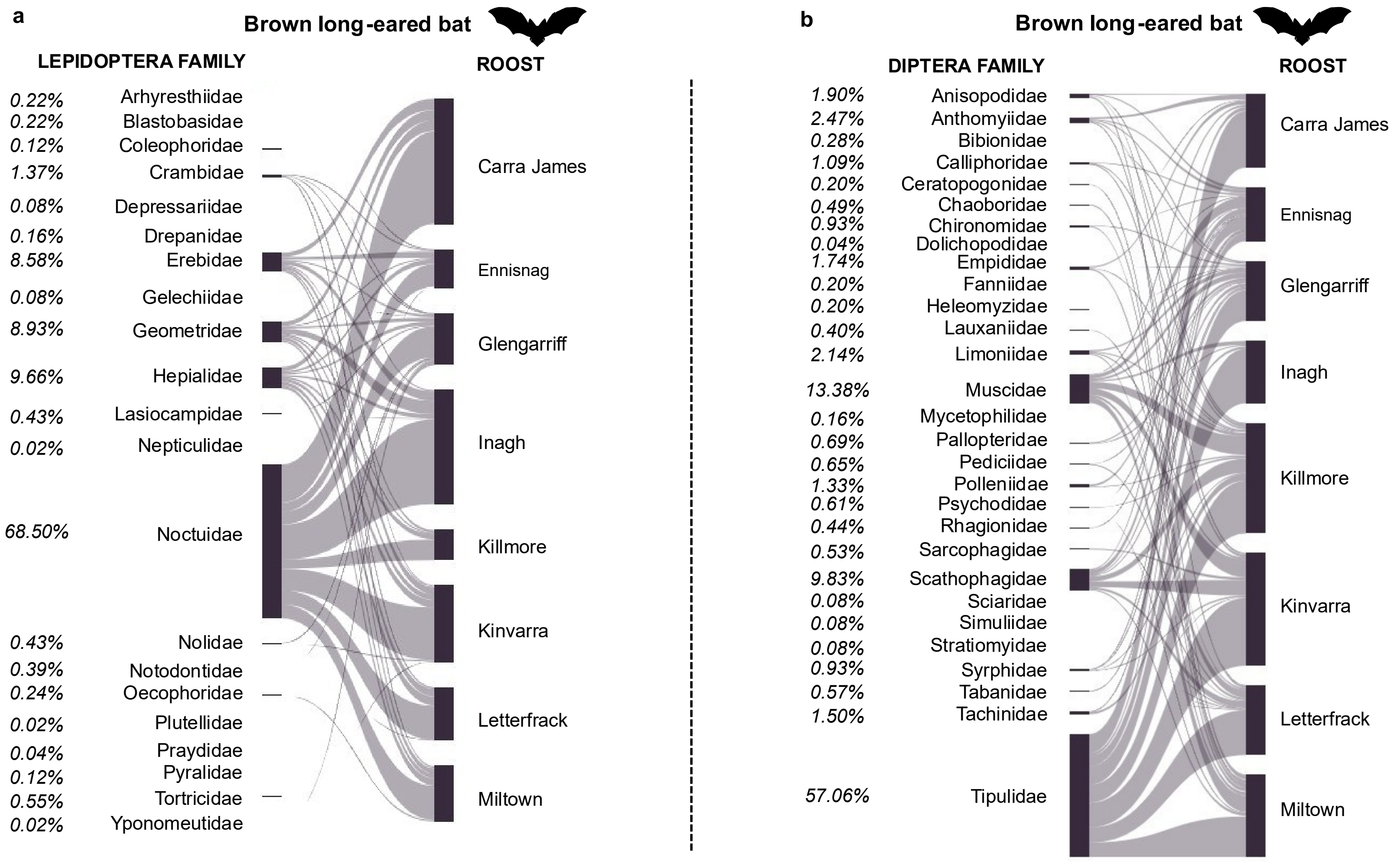


Supplementary Figure S.3A

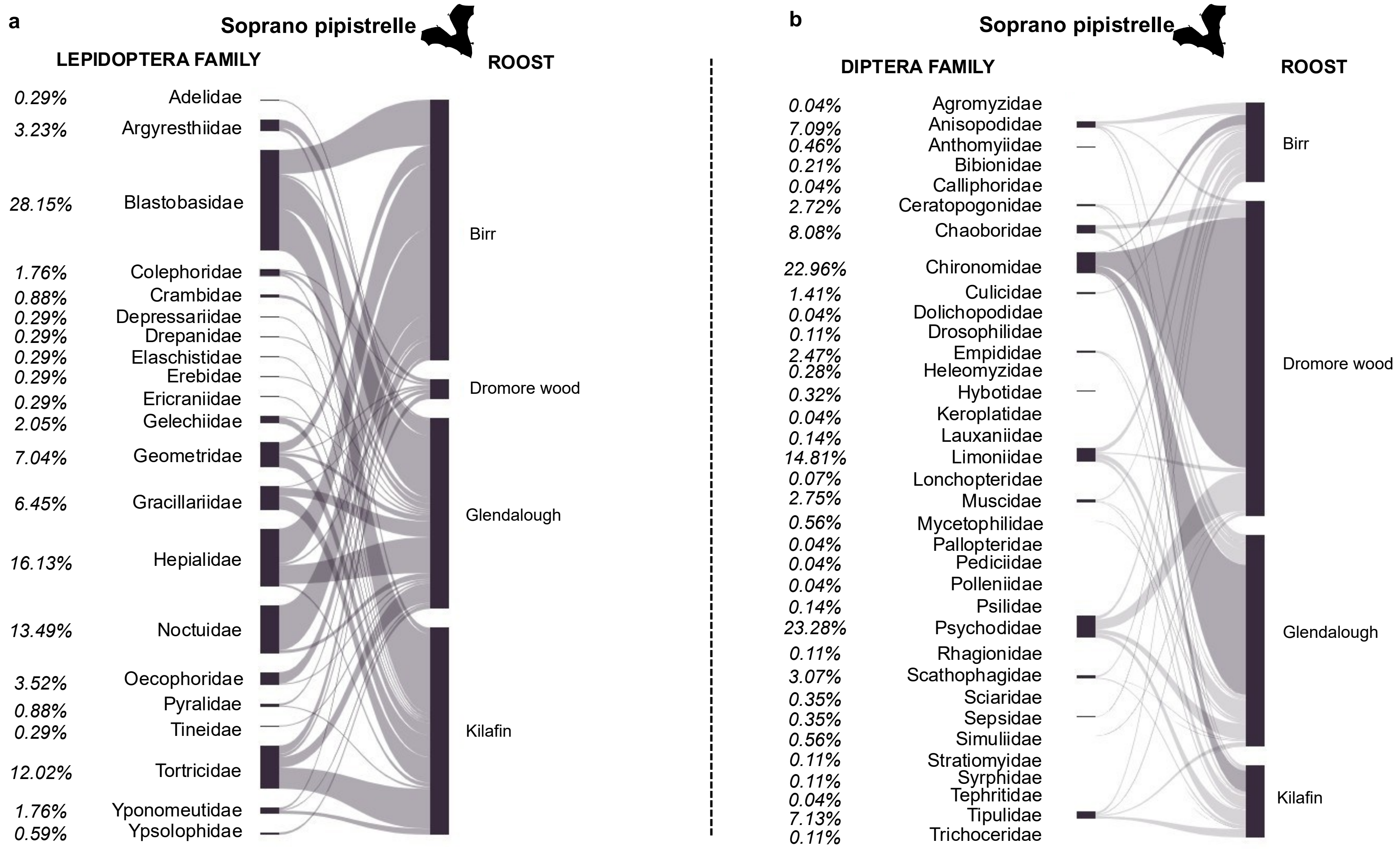


Supplementary Figure S.3B

**Supporting Table S3: Details of outliers discarded for the nMDS analysis.**

| SampleID | Bat species | Roost | Year | Sampling period | nMDS1 | nMDS2 |
|---|---|---|---|---|---|---|
| D1586_21 | Soprano pipistrelle | Glendalough | 2023 | Gestation | -2879.1 | 1153.415 |
| D1631_23 | Brown long-eared bat | Killmore | 2023 | Post-lactation | -10.7066 | -1.50344 |
| D3_2 | Brown long-eared bat | Killmore | 2021 | Gestation | -2.88731 | -2.34257 |
| D1426_13 | Brown long-eared bat | Killmore | 2022 | Post-lactation | -0.34752 | -1.21338 |
| D1426_12 | Brown long-eared bat | Killmore | 2022 | Post-lactation | -0.31472 | -1.21219 |
| D1118_14 | Brown long-eared bat | Glengarriff | 2022 | Gestation | -0.28079 | -1.22061 |
| D1155_5 | Soprano pipistrelle | Kilafin | 2022 | Lactation | -0.02778 | -1.21098 |
| D589_14 | Brown long-eared bat | Killmore | 2021 | Post-lactation | -0.27187 | -1.2112 |
| D589_16 | Brown long-eared bat | Killmore | 2021 | Post-lactation | -0.26608 | -1.2135 |
| D589_6 | Brown long-eared bat | Killmore | 2021 | Post-lactation | -0.26603 | -1.2135 |
| D51_31 | Brown long-eared bat | Miltown | 2021 | Gestation | -0.26425 | -0.21315 |
| D779_14 | Brown long-eared bat | Letterfrack | 2021 | Post-lactation | 3350.001 | 991.1094 |
| D1625_19 | Brown long-eared bat | Letterfrack | 2023 | Post-lactation | -0.02771 | -1.5773 |

**Supporting Information Figure S4**

**Non-metric multidimensional scaling plot showing the dissimilarity in diet composition across bat species (Top) and Brown long-eared with all samples included (Bottom).**

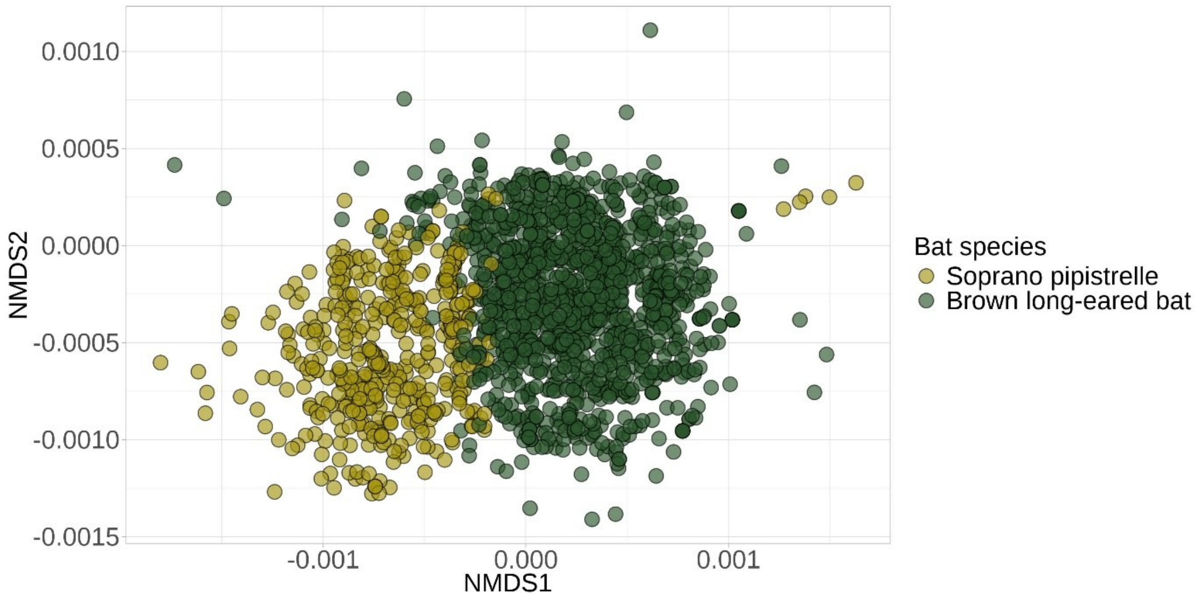


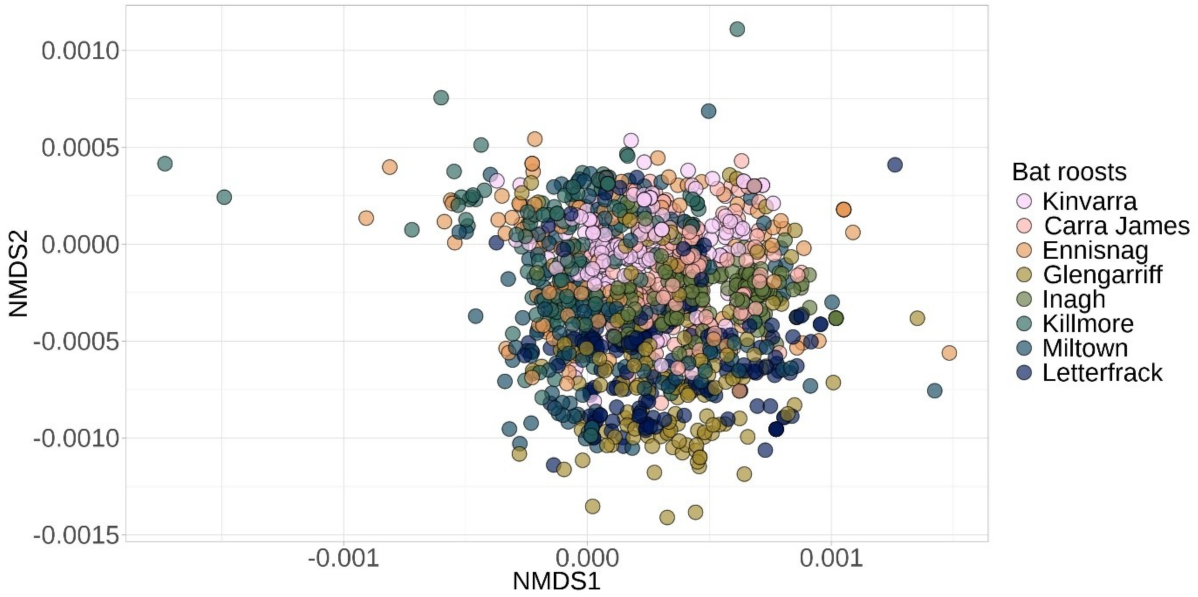
